# Molecular logics in dual sensor regulation of enzyme activity – Phosphorylation OR blue-light activation of cyanobacterial diguanylate cyclases

**DOI:** 10.64898/2026.02.07.704614

**Authors:** Maximilian Fuchs, Andreas Winkler

## Abstract

Bacterial cells use multiple environmental cues to regulate levels of the second messenger cyclic dimeric GMP. This compound influences key lifestyle decisions such as motility, biofilm formation, or virulence. Although many diguanylate cyclases (DGCs) combined with various sensory domains have been studied previously, how distinct inputs are integrated within a single enzyme remains incompletely understood. Here, we investigate a cyanobacterial family of dual-sensor DGCs that combine an N-terminal receiver (Rec) domain followed by a light-oxygen-voltage (LOV) domain upstream of a diguanylate cyclase (GGDEF) domain. Using in vivo activity screening and in vitro characterisation, we determined how phosphorylation and blue light, individually and jointly, regulate enzyme activity. By measuring kinetic parameters across four defined functional states, unphosphorylated or phosphorylated, in combination with dark or light states, we reveal logic gate-like behaviours. One representative, *La*RldC, integrates both signals with pronounced fold-changes in activity-, consistent with overall OR-type logic and with light acting as the dominant input. Our results demonstrate its function as a molecular gate coupling phosphorylation and illumination sensing to cyclic-di-GMP formation. These findings provide valuable insights into multi-signal decision-making in cyanobacteria and establish further understanding of how modular sensory domains are wired to control bacterial second-messenger signalling.

## Introduction

Across all kingdoms of life, cells respond to changing environments by processing diverse physical and chemical cues into behavioural decisions. Among others, phosphorylation, redox state, pH or light conditions can be sensed by a range of modular protein domains [1–5]. The interactions and cross-talk of sensory modules with so called effector or output domains eventually enables intricate molecular mechanisms for multi-signal integration and precisely adjusted physiological responses [6– 8].

In bacteria, the nucleotide second messenger 3’-5’-cyclic-dimeric-GMP (c-di-GMP) orchestrates transitions between motile and sessile lifestyles, biofilm formation, virulence, and cell cycle control [9–15]. Because cellular outcomes depend on c-di-GMP concentration, the enzymes that synthesise and degrade this nucleotide are at the heart of cellular decision-making. The formation of c-di-GMP from two molecules of guanosine-5’-triphosphate (GTP) is catalysed by diguanylate cyclases (DGCs) belonging to the family of GGDEF proteins [16, 17]. Due to their central role for bacterial lifestyle decisions, a range of sensory domains have been fused to these DGCs during evolution, making them versatile natural integrators of environmental inputs. Many GGDEF enzymes carry multiple sensory modules, implying modular logic for decoding diverse stimuli. Exactly how sensory domains effectively couple to the GGDEF catalytic core and tune enzymatic parameters or the oligomeric state, however, remains incompletely understood, particularly in multi-sensor architectures.

Responding to a range of different signal inputs is especially important for cyanobacteria as this lineage experiences pronounced diel transitions due to photosynthesis–respiration cycles that result in oscillations in intracellular pH and redox state [18–20]. These variables, together with changing light conditions, compose a characteristic input landscape that is expected to also impinge on the activity of the frequently occurring DGCs in cyanobacteria. Indeed, cyanobacteria feature a range of light-regulated GGDEF architectures [15, 21–23]. In this study, we characterise a small subgroup of dual sensor DGCs combining an N-terminal Receiver (Rec) domain and a central sensor of Light-Oxygen-Voltage (LOV) domain; a combination that, to our current knowledge, is confined to cyanobacteria. Such Rec-LOV-DGC systems (termed RldCs from here on) might serve as interesting examples of how molecular logics can be realised on a single polypeptide chain.

Rec and LOV domains are frequently occurring sensory systems that respond to phosphorylation and blue light, respectively. Rec domains typically are part of two component systems [24], where they get phosphorylated on a conserved aspartate residue by cognate histidine kinases. In turn, they fulfil their action as response regulator and initiate cellular responses by, for example, tuning transcription factor binding to target gene promoter sequences [25]. In the case of RldCs, the N-terminal positioning of the Rec domain to another sensory element, the LOV domain, raises interesting questions as to how Rec activation would influence blue light processing in LOV. Typically, the flavin cofactor bound to LOV domains is activated by blue light and forms a characteristic photoadduct with a nearby cysteine residue in the cofactor binding pocket [5, 26]. Ultimately, these local rearrangements trigger conformational changes that can be transmitted to a variety of downstream effector domains [27], frequently via helical connectors [28–30]. In cyanobacteria, light strongly impacts pH and the redox state of the cell; thereby illumination might even result in extended multimodal regulation regimes via indirect effects including protonation of residues and/or thiol chemistry.

Interestingly, RldCs feature linker sequences between their sensory modules and the GGDEF domain that suggest parallel coiled-coils as connecting elements (Fig. 1A). Such dimeric helical structures are frequently observed in systems allowing allosteric signal integration (signalling helices, coiled-coils). Especially in the context of GGDEF domains, characteristic register switching mechanisms have been shown to play a central role in signal processing and transduction [28, 31–33]. In the former two systems, either blue light or phosphorylation (LadC or DgcR, respectively) controls GGDEF activity via exactly this mechanism. How light and phosphorylation together can control GGDEF activation, however, remains unresolved and would be interesting to compare to previously described single input systems. Understanding molecular mechanisms of the LOV domain as central hub for signal integration, will also have far reaching consequences for other systems featuring multi domain architectures in combination with LOV domains: for example phototropins [34] or PAS-LOV systems [35].

**Figure 1:**
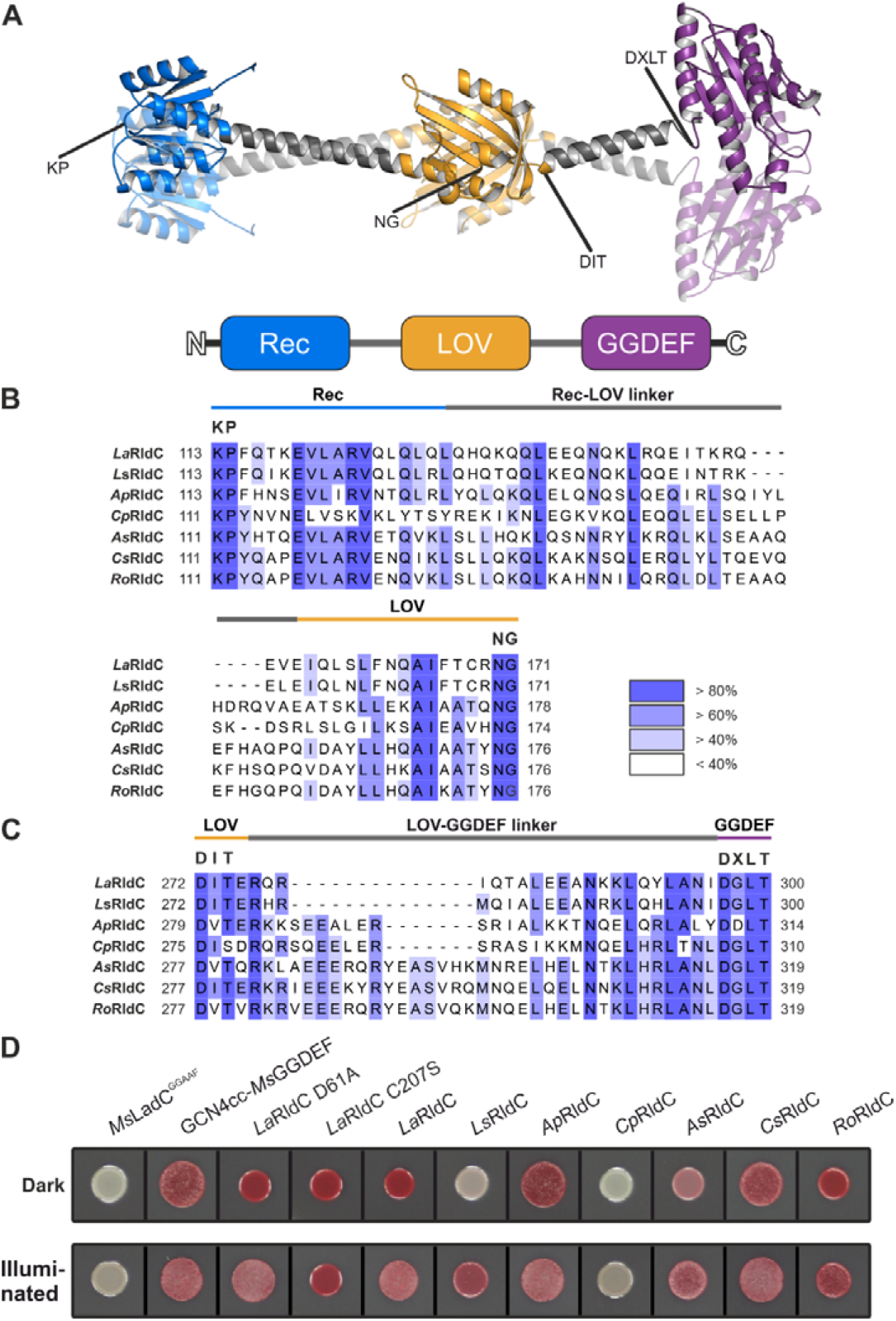
Domain architecture, linker sequence alignments and in vivo activity assay of RldC homologs. A) AlphaFold 2 model of *La*RldC, coloured according to individual domains: Rec – blue, LOV – orange, GGDEF - violet. Linker elements between Rec and LOV as well as LOV and GGDEF are coloured in grey. The individual sequence tags shown in panel B are highlighted. B) Multiple sequence alignment of linker sequences between the Rec and LOV domain of selected homologs. The full sequence alignment is shown in S.1. The KP motif in the Rec domain marks the start of the last loop before Rec-α5. NG marks the conserved start of the first beta strand (Aβ) following the A’α helix of the LOV domain. The legend shows colour coding based on percentage identity. C) Multiple sequence alignment highlighting the three groups with different linker lengths between LOV and GGDEF domain. The LOV-DIT and GGDEF-DXLT motifs flank the linker region. D) The in vivo activity of RldC homologs was investigated using a cellular c-di-GMP level assay. The rdar (red, dry, rough) morphotype is characteristic for elevated c-di-GMP levels. *Ms*LadC^GGAAF^ and GCN4cc-*Ms*GGDEF act as negative and positive controls [31], respectively. Uncut image shown in S.14.

In order to provide a testable framework for quantitative analysis of signal integration, we expressed, purified and biochemically characterized selected representatives of the RldC protein family. In order to describe the cross-talk between individual inputs, we quantified kinetic parameters of several homologs in all four functionally relevant states: Rec unphosphorylated vs phosphorylated in combination with LOV dark vs blue light. Based on previous observations of Rec or LOV activating GGDEF domains, the thereby testable hypotheses would likely be either OR (phosphorylation or light being sufficient to activate DGC activity) or AND gate functionality (LOV activation could act as an enabling second input to phosphorylation – or the other way around). In combination with a mutational inactivation of Rec (phosphorylatable Asp to Ala substitution) and LOV (FMN adduct forming Cys to Ala substitution) we assign input causalities for each domain and their interplay in the context of logic gate functionality. All together, we investigate dual-sensor DGCs as molecular logic devices in which phosphorylation and light inputs are integrated via allosteric coupling to control c-di-GMP synthesis. Among the tested proteins, we identify one promising target for future more detailed studies to fill the gap of missing quantitative rules for signal integration in these dual sensor systems and to provide insights into the molecular logics of Rec-LOV-DGCs. Our study also touches upon rules by which phosphorylation and light might integrate with pH and redox changes, which due to the cyanobacterial origin of RldCs might be of additional relevance.

Defining integration rules in dual-sensor GGDEF enzymes advances our understanding of c-di-GMP signalling architectures that enable nuanced environmental decision-making. These principles can be applied to engineer light- and phosphorylation-gated cyclases, offering precise control over bacterial behaviours and potentially novel optogenetic tools. Understanding how phosphorylation of the Rec domain influences light control of c-di-GMP synthesis, could be employed to generate logic gates that can additionally integrate chemical cues allowing a fine tuning of cellular effects. Conceptually, such system may realize AND, OR, or even antagonistic (‘NAND’ or ‘NOR’) logic depending on the sign and magnitude of the coupling between Rec, LOV and the target output domain. As far as significance and broader impact are concerned, understanding dual-input integration has implications for bacterial decision-making processes, spatial and temporal control of the bacterial second messenger c-di-GMP, and future synthetic biology/optogenetics approaches.

## Results

Using InterPro and PSI-BLAST, we gathered an initial set of approximately 1,500 sequences featuring a Rec-PAS-GGDEF architecture. We filtered these by looking specifically for Rec-LOV-GGDEF architectures, ensuring the presence of specific conserved residues for individual domains: Rec – conserved aspartate as phosphorylation site, LOV – GX(N/D)C(R/H)(F/I)L(Q/A) motif containing the adduct-forming cysteine, GGDEF – GG(D/E)EF motif. The largest number of sequences was removed by filtering for the LOV motif; only around 10 % of the sequences contained the canonical motif and were considered to act as blue-light receptors. A few sequences had to be removed since they included additional PAS or EAL domains, highlighting that even more complex domain architectures can be found in nature. Eventually, the reduced alignment of RldC homologs satisfying all constraints retained 15 sequences (S.1). Using AlphaFold2 predictions, we identified helical linkers connecting individual protein domains (Fig. 1A). The linker between Rec and LOV domain extends the α5 helix of the Rec domain and connects it to the A’α helix of the LOV domain [27, 36]. Fig. 1B shows a sequence alignment starting at the conserved KP motif in the Rec domain and ending at two conserved residues placed at the beginning of the first beta-strand (Aβ) of the LOV domain. We identified characteristic linker length differences with 7-residue offsets, as frequently observed in coiled-coil linkers and their heptad repeats [37, 38]. The linker between the LOV and GGDEF domains also shows similar variability (Fig. 1C), with length differences of one or two heptad repeats. Since different linker lengths have previously been shown to impact fold-changes of enzymatic activity in diguanylate cyclase effectors [28] we selected two homologs per linker length for a more detailed characterisation.

### In vivo screening

After cloning the genes into a pET-M11 expression vector, we performed an initial in vivo DGC activity screening: one, to assess the overall activity, and two, to estimate the influence of blue light. The screening is based on the production of cellulose and amyloids due to increased c-di-GMP levels in *E. coli*. We were able to utilise the leaky expression of the Lac operon, since the activity of most homologs is apparently relatively high. The Congo red dye incorporated into the culture media accumulates at cellulose and amyloid fibres, resulting in a distinct visual appearance due to the so-called rdar (red, dry, rough) morphotype [39]. While the assay can be used to address overall diguanylate cyclase activity as well as up-or downregulation caused by blue light, the influence of different expression levels or proper folding cannot be easily assessed. However, as an initial qualitative analysis, it can be a valuable approach for mid-to high-throughput screening. In addition to positive and negative controls from an established system [31], we included additional controls in the screening to better interpret morphological changes. The variant *La*RldC D61A removes the Asp that would typically be phosphorylated, while C207S removes the Cys residue necessary for the protein’s photocycle. The screening results showed that the morphotype of individual homologs varies significantly [Fig. 1D]. We categorised them based on the observed effects. 1 – Promising: *As*RldC, *La*RldC, and *Ls*RldC. While basal activity varied substantially, we observed pronounced differences between dark and illuminated plates. 2 – Neutral: *Ap*RldC, *Cs*RldC and *Ro*RldC. All showed diguanylate cyclase activity, but no apparent blue light regulation. However, for *Ap*RldC and *Cs*RldC, high expression levels could mask a light response. 3 – No signal: *Cp*RldC. The morphotype of this homolog resembled the negative control. While this could mean that the protein is not producing any c-di-GMP, we cannot exclude other factors linked to the previously mentioned drawbacks of the screening. Since the assignment of individual homologs to these categories did not correlate with linker length variations, we proceeded to expression and purification for a more detailed in vitro characterisation.

### Protein production

We produced all homologs using the corresponding codon optimised coding sequences cloned into the pET-M11 vector and *E. coli* BL21 (DE3). Although all genes were overexpressed in *E. coli* (S.2), we were unable to identify suitable conditions for extracting and purifying *As*RldC and *Cs*RldC. This was most likely due to the formation of inclusion bodies. Considering the complex domain architecture and the successful purification of other homologs, additional refolding attempts were not pursued and, instead, we focused on the characterisation of the proteins obtainable in soluble form.

### Protein purification

We were able to purify all remaining homologs with acceptable yields. Intact and native MS measurements were used to confirm molecular weight, oligomeric state, and cofactor binding (S. Table 1). During protein quantification, we observed a shoulder to the 280 nm peak for *Ap*RldC, *La*RldC and *Ro*RldC, likely due to bound c-di-GMP (S.3A). The spectrum of guanosine has a maximum at 254 nm and would therefore fit to the observed shoulder. For *Ro*RldC specifically, the elution volume during size exclusion chromatography was lower than that of the other homologs, which suggests that c-di-GMP binding led to a higher oligomeric state. However, native MS measurements did not show clear indications of higher oligomers. *Cp*RldC and *Ls*RldC did not show signs of bound c-di-GMP (S.3A). Both homologs showed low enzymatic activity in the in vivo screening, which suggests that cellular c-di-GMP levels are lower during protein production. Although *Ls*RldC has high sequence similarity to *La*RldC, its purification proved to be more challenging, primarily due to protein aggregation. To determine whether bound c-di-GMP could enhance protein solubility, we examined protein production under constant blue light illumination, which was expected to increase cellular c-di-GMP levels. This procedure increased yield, but solubility remained an issue even when c-di-GMP was bound. While bound c-di-GMP should not influence the spectroscopic characterisation, it is problematic for kinetic measurements, since it affects overall enzymatic activity and might affect fold changes between dark and light state [40].

### c-di-GMP removal and disulfide reduction

To remove bound c-di-GMP, we used the phosphodiesterase RocR [41]. In the case of *Ap*RldC, removal of c-di-GMP led to protein aggregation and eventually precipitation. Although multiple conditions were tested, the remaining soluble protein still contained bound c-di-GMP. For *La*RldC, increasing the ionic strength of the buffer mitigated the decreased solubility. Upon removal of c-di-GMP, however, intact mass measurements showed the formation of a disulfide bond between the two protomers (S.4A). To successfully prevent this oxidation process (S.4B), we supplemented the corresponding buffers with TCEP (S. Table 2). *Ro*RldC also showed decreased solubility after c-di-GMP removal. Increasing the buffer’s salt concentration improved solubility. Additionally, we observed an increased elution volume during SEC compared to before, indicating a shift toward the dimeric species (S.5). While complete removal of c-di-GMP was confirmed via HPLC (S.6), we also observed characteristic spectral changes in the UV region of the absorption spectra in the product free protein stocks (S.3B). Extending the spectroscopic characterisation to the Vis-region, the characteristic features of the LOV domains are apparent for all well-behaved systems (Fig. 2A - E).

**Figure 2:**
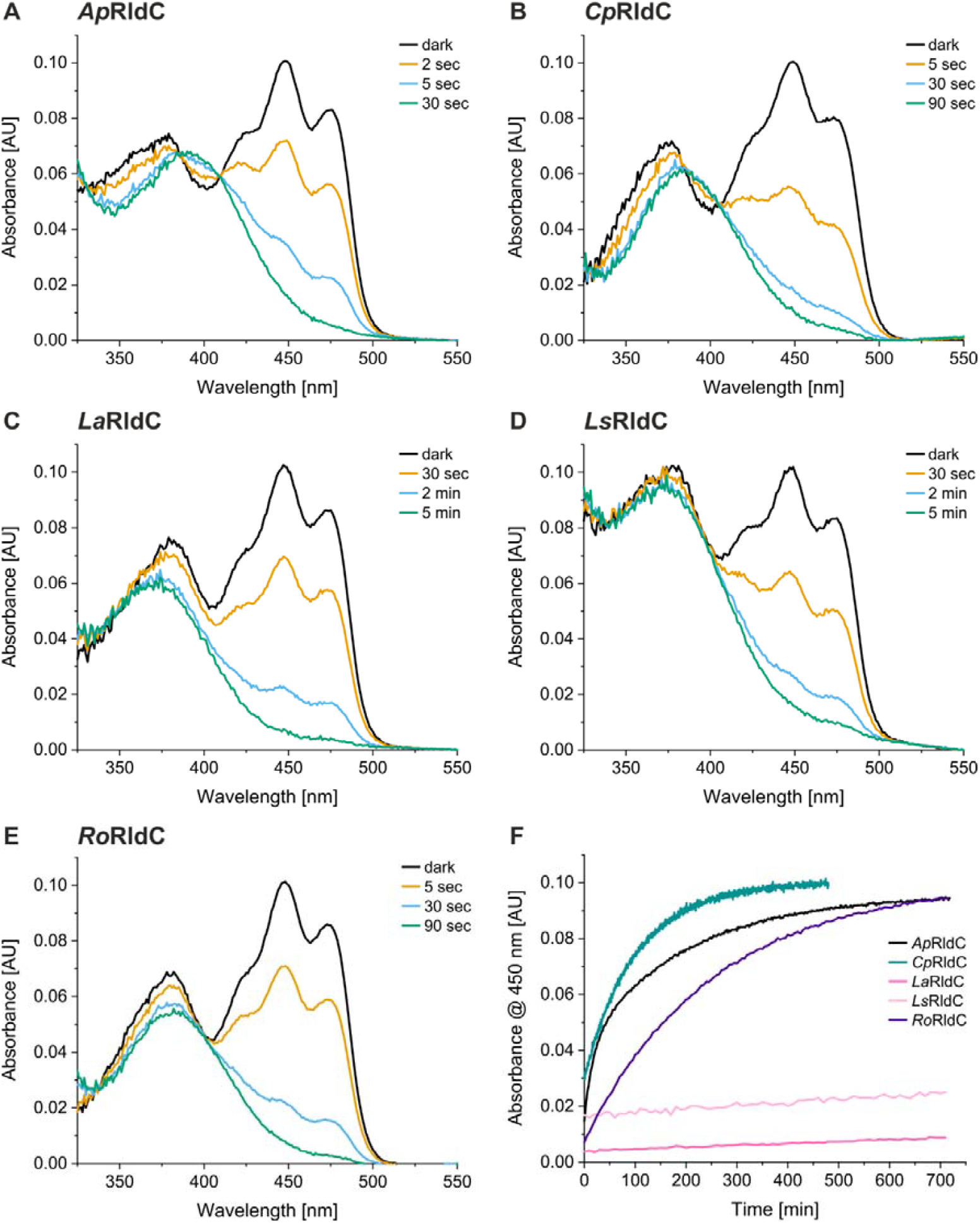
Blue light response and dark state recovery of RldC homologs. A) *Ap*RldC shows canonical spectral changes and fast adduct formation. B) *Cp*RldC, which also features canonical spectral changes, but slower adduct formation compared to *Ap*. C) *La*RldC with atypical light state spectrum. D) *Ls*RldC similar to *La*RldC. Spectrum is influenced by impurities and its tendency to aggregate. E) *Ro*RldC also shows atypical light response but with faster activation compared to *La*RldC and *Ls*RldC. F) Dark state recovery of all characterised homologs. The 450 nm absorbance is used to follow the thermal relaxation of the photo-adduct. Activation and thermal relaxation were measured at 20 °C.

### Spectroscopic characterisation

The illumination of LOV domain-containing photoreceptors with blue light leads to characteristic spectral changes caused by the adduct formation between a conserved cysteine residue and the C4α of the isoalloxazine ring of the FMN cofactor. *Ap*RldC and *Cp*RldC showed LOV characteristic spectral changes (Fig. 2A, B). Upon illumination, the 370 nm band red-shifts to 390 nm, and the 450 nm absorption decreases. The other three homologs show what one might consider an atypical response to blue light (Fig. 2C, D, E): the 370 nm band does not show the characteristic red-shift. This is particularly pronounced for *La*RldC and *Ls*RldC. This shifts the two isosbestic points, usually observed for the dark and light states, to lower wavelengths for *Ls*RldC and *Ro*RldC. In the case of *La*RldC, only one isosbestic point remains. During attempts to identify the molecular origin of this unexpected behavior, we observed that the wavelength of the light source has a significant influence on the percentage of photoproduct (S.7), i.e. the photostationary state (PSS) [42, 43]. Two of the atypical photoproduct homologs, *La*RldC and *Ls*RldC, could only be fully activated using a 470 nm LED (Fig. 2C, D), but not a typically employed 455 nm light source. This might be caused by a more significant overlap in the near UV region of the 455 nm LED, which may partially induce photoconversion back to the dark state, leading to a steady state of the two populations. *Ro*RldC, another homolog with a similar behaviour regarding the UVA spectral region, also benefited from using the 470 nm LED, but with less significant differences between the two light sources.

In addition to reverse photoactivation, LOV domains also thermally revert back to their ground state. We recorded dark recovery data for all homologs (Fig. 2F) and generally observed slow thermal relaxation rates. The mean lifetimes were estimated using single-exponential fits (S.8). *Cp*RldC has a mean lifetime of ∼100 min and is the fastest recovering homolog. Due to its tendency to aggregate upon prolonged incubations, the corresponding trace in Fig. 2F is only shown for the initial part of the measurement. Other homologs recover somewhat slower; *Ap*RldCs mean lifetime is 150 min, and *Ro*RldCs 254 min. *La*RldC and *Ls*RldC deviate substantially and show particularly slow recoveries, resulting in very stable light states, preventing a fit to the dark state recovery kinetics for these two homologs. However, we demonstrated that illumination with UV light (395 nm LED) results in efficient reversion to the dark-adapted state (S.9).

The observed atypical spectral changes raised the question of whether the affected homologs form the characteristic LOV flavin-cysteinyl adducts. Therefore, we used the previously created C270S variant of *La*RldC, which substitutes the adduct-forming cysteine, to check its response to illumination in vitro. The variant did not respond to blue light at comparable intensities to those used for the wild-type protein (Fig. 3A). Even illumination with maximum intensity only moderately influenced the absorption spectrum of the cofactor; actually resulting in cofactor bleaching and no recovery to the starting spectrum in the dark. In addition, we observed a broader 370 nm peak for the C207S variant compared to the wild type, but reminiscent of the supposed photoadduct signatures of *Ls*RldC and *La*RldC. To investigate if atypical photoadduct spectra could result from cofactor reduction, we performed a photoreduction under anaerobic conditions [Fig. 3B]. This showed that the light response of the wild type is different from the spectral changes caused by the reduction of the cofactor. The photoreduction was slow and did not proceed via semiquinone formation. Importantly, the recovery of the starting spectrum via reoxidation of the cofactor was significantly faster than the recovery of the photoadduct. Although the spectral properties of some homologs are atypical, the results confirm the functionality of the LOV domain as a blue light photoreceptor.

**Figure 3:**
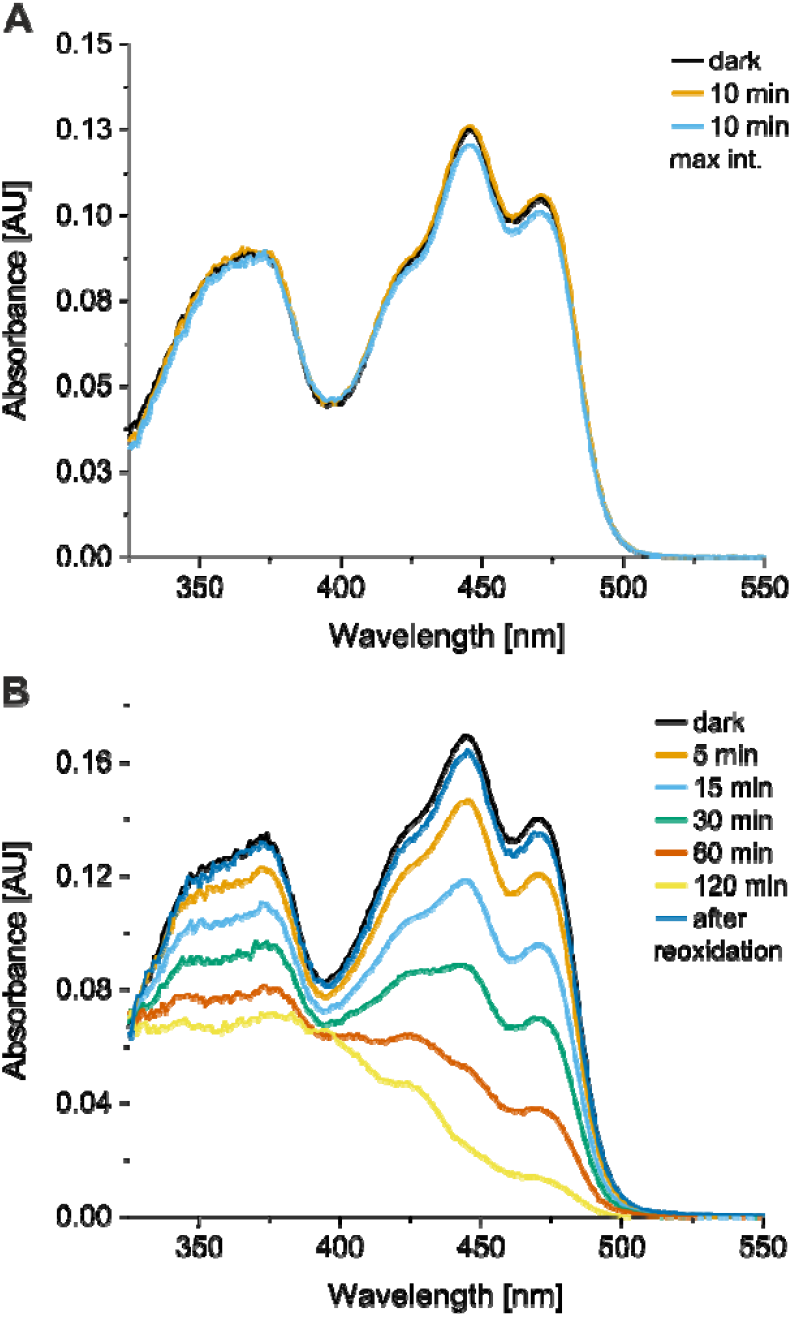
Characterisation of LaRldC C207S. A) Absorption spectrum of *La*RldC C207S in response to illumination. B) Photoreduction of the C207S variant using EDTA as an electron donor, combined with methyl viologen and 5-deaza-FAD, to increase electron transfer efficiency. FMN reoxidation, after opening the sealed quartz cuvette is also shown.

### Phosphorylation experiments

For the second sensory domain, the Rec domain, phosphorylation is typically mediated by cognate histidine kinases, which autophosphorylate in response to diverse signals. The phosphoryl group can then be transferred to the conserved aspartate of the respective Rec domains. While this process is usually the most efficient way to phosphorylate a Rec domain [44], small-molecule phosphodonors such as acetylphosphate and phosphoramidate (PA) can also be employed. We used a Phos-Tag™ SDS-PAGE to separate and detect unphosphorylated and phosphorylated proteins. The migration in these gels is usually slowed down in comparison to a normal SDS-PAGE and although the homologs have similar masses, the distance between *Cp*RldC and the other homologs bands is quite large (Fig. 4A). However, we previously confirmed the masses using intact mass measurements (S. Table 1), suggesting that Phos-Tag™ gel migration behaviour is indeed difficult to predict. A comparison of the bands showed that *La*RldC and *Ro*RldC can be successfully phosphorylated using ammonium hydrogen phosphoramidate (Fig. 4A). However, the degree of phosphorylation varies among homologs. For *La*RldC, phosphorylation appears to be pH-dependent, with a more neutral pH increasing the amount of phosphorylated protein. *Ro*RldC was only partially phosphorylated under the tested conditions, while *Cp*RldC is not readily phosphorylated at all. *Cp*RldCs features an amino acid substitution D16K, which has previously been shown to reduce phosphorylation efficiency [45]. While extended incubation times allowed us to detect phosphorylated protein during intact mass measurements (S.10) the low ratio between phosphorylated and unphosphorylated protein remained a hindrance to further exploring the effects of phosphorylation on enzymatic activity or other functional properties in this specific case.

**Figure 4:**
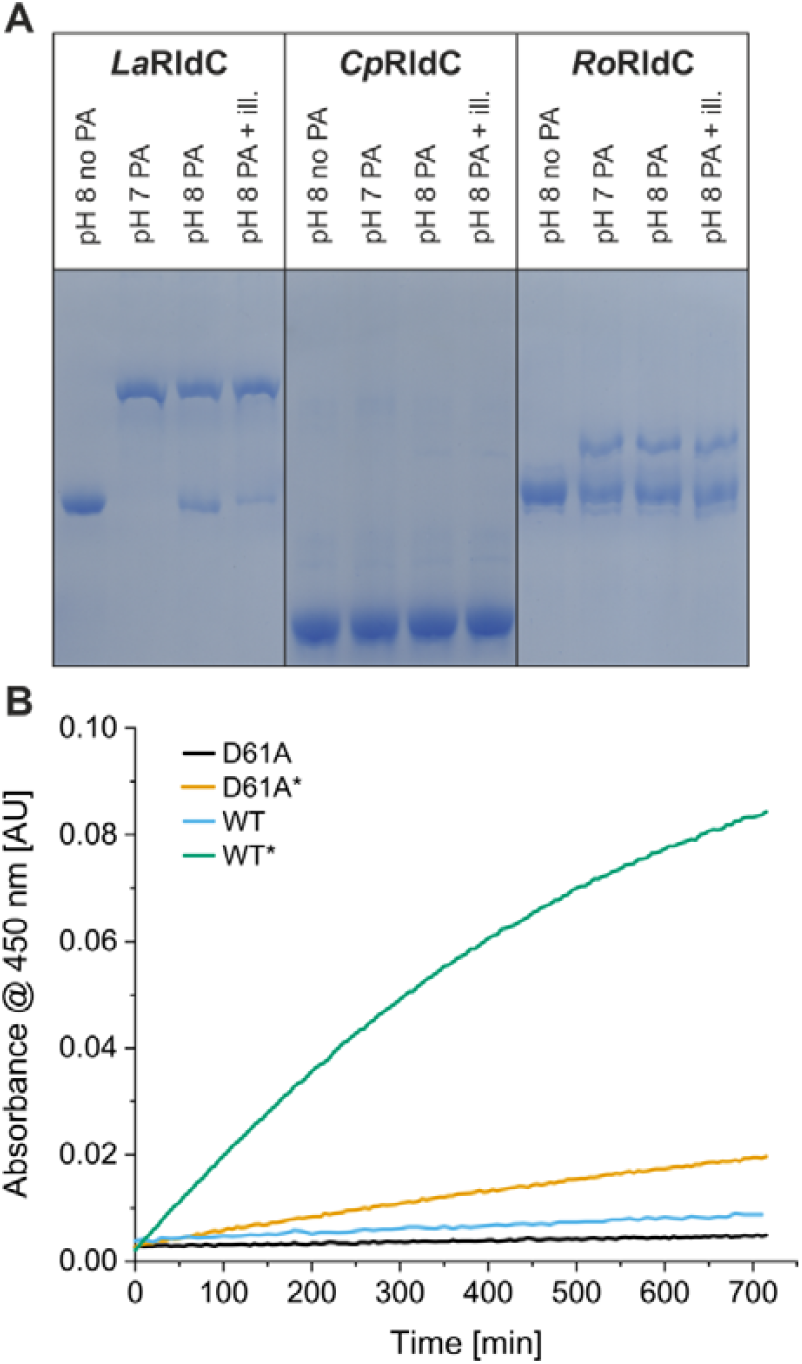
Rec domain phosphorylation and implications for LOV photocycle properties. A) Phos-Tag™ SDS PAGE analysis of the amount of phosphorylation after 1 hour of incubation with 50 mM PA under different conditions. Unless indicated, samples were kept in the dark state during incubation. Complete SDS-PAGE image shown in S.15. B) Changes in dark state recovery of *La*RldC WT and the D61A variant before and after incubation with 50 mM PA as indicated by an asterisk.

Since the Rec domain precedes the LOV domain, we also addressed the effect of phosphorylation on the photocycle. Indeed, we observed pronounced changes in the recovery kinetics of *La*RldC (Fig. 4B). The phosphorylation of *La*RldC effectively increases the protein’s recovery rate constant. While the unphosphorylated protein barely reverts to the dark state, phosphorylation drastically increases the recovery kinetics. This could be partially attributed to changes in ionic strength and the presence of additional ions resulting from the dissolved ammonium hydrogen phosphoramidate, which is necessary to maintain high levels of phosphorylation. However, control experiments with an unphosphorylatable D61A variant of *La*RldC showed that the effect on the recovery rate is mainly due to phosphorylation. For *Ro*RldC, we also observed an increased recovery rate constant, but to a lesser extent, potentially due to a lower fraction of phosphorylated protein and to changes in ionic strength. Interestingly, the blue-light activation kinetics of *La*RldC were not affected by protein phosphorylation (S.11). While the initial set of homologs was reduced to three, we still had one member of each LOV-GGDEF linker length group moving forward to assess diguanylate cyclase activity in vitro.

### In vitro activity

To investigate the in vitro activity of *La*RldC, *Cp*RldC and *Ro*RldC, we measured initial velocities (v0) of the homologs using an offline HPLC-based assay. For initial comparisons, all in vitro activities were determined at a GTP concentration of 100 µM (Fig. 5), since other GGDEF-proteins have typically been characterised using GTP concentrations between 50 to 500 µM [28, 31–33, 46]. To avoid any impact of high PA concentrations on activity, we diluted the samples directly before starting the reaction, thereby reducing the PA concentration from 50 mM to 5 mM. All homologs showed some degree of light activation, but *La*RldC showed the biggest change with a 73-fold increase. Due to the high activity of the protein, we could only use the 5 and 10 second timepoints to determine v0. Reducing the enzyme concentration further was not an option, as concentrations below 1 µM decreased specific activity, most likely due to a monomer-dimer equilibrium (S.12). *Cp*RldC and *Ro*RldC exhibited smaller changes of 2-fold and 2.5-fold, respectively. In line with the in vivo screening, *Cp*RldC had a significantly lower specific activity than the other two homologs. In addition, the conversion of GTP was accompanied by the formation of the reaction intermediate 5’-pppGpG, suggesting suboptimal alignment of the GGDEF protomers [47]. Since phosphorylation of *Cp*RldC was not feasible, only *La*RldC and *Ro*RldC were characterised regarding their response to the second input signal. Phosphorylation of dark state *La*RldC increased activity 16-fold; shifting the protein to the light state further increased activity 5-fold. On the other hand, *Ro*RldC increased activity upon phosphorylation in the dark state only by 1.5-fold and a further 2.0-fold increase when illuminated. These results show that the individual signals and the combination of both can modulate DGC activity. However, the extent of the regulation differs substantially between homologs.

**Figure 5:**
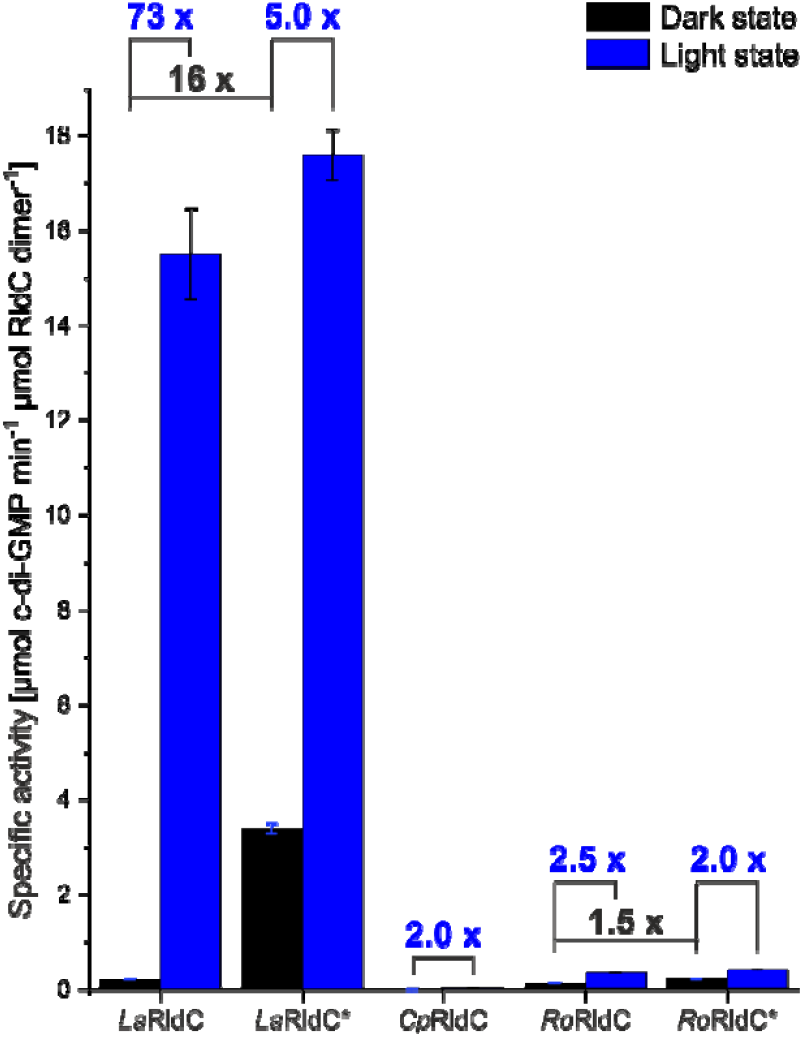
Specific activity of RldC homologs. Values for 50 µM substrate concentration are shown. Dark state activities are depicted in black and light state activities in blue. * Indicates phosphorylated protein. Fold changes between states are indicated above the bars.

## Discussion

Detailed studies of dual sensor photoreceptors remain scarce, most likely due to the complex nature of these proteins and their frequently unknown physiological roles [48, 49]. Here, we characterised several dual-sensor diguanylate cyclase homologs and investigated their responses to both illumination and phosphorylation.

The initial in vivo screening enabled qualitative analysis of the enzymatic activity of all selected homologs under both dark and illuminated conditions. *Cp*RldC appeared inactive on the plate screen (Fig. 1D) and was later shown to exhibit very low specific activity, yet measurable light regulation in our in vitro measurements. Since we investigated multiple homologs, we tried to maintain comparable buffer conditions throughout our analyses. Yet, for some proteins, these conditions might not be optimal, and additional studies could be performed to improve activity by varying pH, temperature, and/or ionic strength [50]. The variants of *La*RldC exchanging the phosphorylated aspartate for alanine (D61A) and the adduct-forming cysteine for serine (C207S) were used in the in vivo assay as additional controls. For one, it checks for any cross-talk of histidine kinases present in *E. coli* Bl21 (DE3) with the Rec domain. On the other hand, it helps to confirm that the sometimes-subtle differences between dark and illuminated plates are due to light-induced changes in enzymatic activity. While we identified several homologs that showed light regulation in vivo, not all could be purified due to protein solubility issues. Interestingly, due to their high DGC activity, some homologs were copurified with c-di-GMP, most likely bound to the I-site of the GGDEF domain [40], which is conserved in all investigated homologs. When we performed native MS measurements to confirm cofactor loading and the dimeric state of all homologs, however, we did not observe evidence of bound c-di-GMP. This suggests that the interaction between product and protein is not strong enough to withstand ionisation or is lost during the exchange from storage buffer to ammonium acetate. The feedback inhibition via the I-site might also complicate the interpretation of the in vivo screening, since it could affect the homologs to different degrees. To mitigate such drawbacks, we proceeded to in vitro assays and used RocR [41] as an established EAL phosphodiesterase to successfully remove c-di-GMP from *La*RldC and *Ro*RldC. Interestingly, product inhibition at the I-site substantially influences conformational dynamics, as in the case of *La*RldC, a disulfide bond between the two protomers formed rapidly after c-di-GMP removal. Disulfide bond formation was likely due to a cysteine residue in the A’α helix of the LOV domain. The residue is part of the coiled-coil linker connecting the Rec and LOV domains, which probably influences communication between the two. In fact, this might suggest additional cross-talk between the LOV domain, tentative signal integration pathways from the Rec domain and the effector domain with intracellular redox state. We only observed this disulfide bond in *La*RldC and *Ls*RldC, where this specific cysteine is conserved. Structure predictions of the dimers also showed the close proximity of these cysteines. In kinetic measurements (data not shown). Using the oxidised species of *La*RldC, we observed lower substrate turnover and reduced light activation. Potentially, cysteine oxidation could serve as an additional input on RldC regulation as observed in other redox-sensing systems [51–53]. It was also reported that cyanobacteria exhibit a certain light/dark modulation of thiol oxidation [19]. Therefore, the additional influence of light on pH and redox might even result in extended multimodal regulation regimes. As pH can alter protonation states of catalytic residues and metal coordination at the Rec and GGDEF domains, it might further influence RldC activity; along this line, cellular redox can also influence the LOV cofactor redox state and/or cysteine thiol chemistry. Eventually, all of these mechanisms might be used to tune allosteric communication. Thus, Rec phosphorylation and LOV photoactivation operate within a broader, time-varying chemical environment that controls GGDEF activation. While these findings leave multiple open questions regarding output regulation, we did not further investigate redox effects at this time and instead focused on illumination and phosphorylation.

To address the effects of illumination, we analysed the spectroscopic properties of all purified RldC homologs. We identified three representatives that did not exhibit the characteristic LOV photoproduct with an absorption maximum at 390 nm. Instead, the absorption maximum was blue-shifted towards 370 nm (S.13A). This unusual behaviour coincided with the necessity to use a 470 nm LED for maximum photoconversion, which was especially apparent for *La*RldC and *Ls*RldC. While influences of light quality have been reported for LOV domains before [54], it is intriguing to observe such a clear preference for higher wavelengths in terms of activation, despite the dark state spectrum not being red-shifted itself (S.13B). Previous studies have shown that ZEITLUPE (ZTL) from *Arabidopsis* also shows a blue-shift of the light-state spectrum compared to other LOV proteins [55]. Interestingly, there appears to be some correlation with a slow thermal reversion of the photoadduct in LOV domains and these peculiar spectral properties. ZTL and *Ro*RldC actually exhibit similarly slow dark state reversions. The even slower recovering *La*/*Ls*RldCs feature an even stronger blue shift of the flavin cysteinyl adduct spectrum. However, there are clearly also other factors influencing dark state recoveries, as very slow LOV proteins, like for example FKF1, can also feature characteristic LOV-adduct spectra [56, 57]. Spectral tuning of LOV domains is considered interesting for multiple applications, including optogenetics. While this usually focuses on changing the properties of the dark state, not the light state, using lower-energy, and therefore less damaging, radiation for activation could be beneficial. We tried to identify the specific residues responsible for the shifted light-state spectrum by examining the cofactor’s surroundings. Therefore, we used the dark-adapted and light-adapted structures of AsLOV2 [58] to identify residues and compared them with those of our purified homologs and ZTL. While some differences were observed for *Cp*RldC near the flavin dimethyl-benzyl ring, this homolog’s spectrum showed no atypical characteristics. For those homologs with a clearly shifted spectrum, like *La*RldC, no obviously deviating amino acids could be identified, suggesting that second shell residues might exert this intriguing influence.

To confirm that we indeed observe photoadduct formation, we also characterised a C207S variant of *La*RldC. While the dark-state spectrum of the protein changed due to the exchange, similar amino acid substitutions have already been shown to alter the absorption spectrum of other LOV proteins slightly [59, 60]. Generally speaking, the flavins’ short wavelength peak depends more on solvent polarity and hydrogen bond formation, and can therefore be influenced more readily [54]. Without the adduct-forming cysteine, we no longer observed a LOV-characteristic response to blue light. Photoreduction of the variant under anaerobic conditions also yielded a qualitatively different spectral change compared to the wild type. Together, these findings support photoadduct formation in all our systems and hint at subtle differences in the local flavin environment influencing the electronic properties of the flavin. Additionally, once exposed to oxygen, the reduced FMN reoxidized swiftly, in contrast to the slow recovery of the very stable light state observed for *La*RldC. These characteristics may hint at the protein’s physiological role. Studies of photoreceptors in *Arabidopsis thaliana* have shown that LOV domains with significantly different recovery rates, namely ZTL and FKF1, are associated with specific functions. FKF1, which slowly reverts to the ground state, operates in a day-loop, whereas the fast-recovering ZTL senses light-dark transitions [56]. While a very stable light state might not be desirable for all optogenetic applications, such a behaviour could prove interesting, as illumination with near-UV light can efficiently shift the protein back to its dark state, thereby eliminating the need for constant illumination while retaining light control. In contrast to what was shown for LOV2 of *Adiantum*, this process is not faster than activation but rather occurs on a similar timescale [6].

To test the effect of phosphorylation, we used the small-molecule phosphodonor phosphoramidate, which features a nitrogen-phosphate bond similar to that of natural activated histidine kinases. The compound, in fact, was able to phosphorylate several RldCs. However, depending on the homolog, the degree of phosphorylation varied substantially and was further influenced by the phosphorylation conditions. In the case of *La*RldC, we found that a lower pH led to more complete phosphorylation. Similar observations were made for a Rec domain that is part of a virulence switch in *Salmonella*. There, pH dependence was attributed to a histidine residue proximal to the phosphorylated aspartate [61]. A sequence alignment revealed that this histidine residue is also present in *La*RldC, and structure predictions indicate its proximity to Asp61. *Cp*RldC and *Ro*RldC do not feature this histidine residue and did not exhibit a clear pH dependence. In addition to the pH dependence of *La*RldC, we observed that illumination (Fig. 4A) increases the amount of phosphorylated protein. This could be caused by light-induced changes at the dimer interface, including the A⍰α helices, that favour or stabilise the phosphorylated state. Since signal transduction from the Rec domain via the LOV domain to the DGC will likely depend on the domain-connecting coiled-coil linkers, such an allosteric cross-talk of the individual sensors should be an integral part of the molecular signalling mechanism. In fact, the influence of substituting the phosphorylatable Asp in the Rec domain (D61A variant) on thermal recovery of the LOV domain further highlights such an allosteric communication pathway between domains; a feature frequently demonstrated for sensory proteins using methods to address their conformational dynamics in various functional states [29, 62–67].

In the case of *Cp*RldC, neither Phos-Tag™ SDS-PAGE nor intact MS measurements showed substantial amounts of the phosphorylated species. An Asp to Lys amino acid exchange of one of the aspartate residues, usually interacting with the essential Mg2+ ion bound by Rec domains, might mitigate the phosphorylation via small molecule phosphonors. Studies investigating CheY, a stand-alone Rec domain, have shown that a similar variant can still be phosphorylated by its cognate kinase CheA, but that the efficiency was lowered [45]. It should be noted that this variant of CheY exhibits similar behaviour to the phosphorylated Rec domain and can therefore be considered a constitutively active variant. However, structural studies showed that wild type and variant adapt the same structure when crystallised [68]. Therefore, we did not expect *Cp*RldC to exhibit a similar behaviour, as the proposed activation mechanism would involve structural rearrangements transmitted through the alpha-helical linkers rather than protein-protein interactions.

Similar to the phosphorylation experiment using Phos-Tag™ gels, preliminary kinetic measurements assessed the effect of pH on enzymatic activity. We observed an influence of buffer pH on the activity and fold changes of the studied homologs. Interestingly, measurements at pH 7.0 yielded lower fold-changes, whereas measurements at pH 8.0 yielded higher fold-changes. While this contrasts the observed phosphorylation efficiency via Phos-Tag™ analysis, we chose to use pH 8.0 for a more detailed kinetic characterisation; also, because cyanobacteria have been shown to have a higher cytosolic pH in light, which assists carbon fixation [20]. Since the in vivo screening reflects the activity and fold changes at the physiological pH of *E. coli* cytosol, it also explains why the in vitro fold change*s* of *La*RldC are much higher than indicated by the screening. The in vitro measurements showed that phosphorylation of the N-terminal Rec domain increases activity by 16-fold, while illumination results in a 73-fold change. Although both individual signals and their combination led to higher activity and hence reflects an OR logic, this might not describe the protein’s behaviour in every detail. Phosphorylation of the Rec domain has less impact than illumination, suggesting that the organism primarily more readily responds to light to upregulate c-di-GMP production and that the signal that triggers phosphorylation has less impact. This provides the opportunity to fine-tune cellular responses in response to external factors. The observed influence of phosphorylation on dark-state recovery and the increased phosphorylation ratio in the light-state add further complexity. These observations highlight the allosteric nature of both signals. While we were able to characterise the effect of phosphorylation for this homolog, it is unclear which external factors control it. The gene preceding *La*RldC appears to encode a hybrid histidine kinase that features an extracellular Cache domain (dCache_1, PF02743). While Cache domains are the most abundant extracellular sensor domains in prokaryotes, their ligands are diverse and still remain largely elusive [69, 70]. Therefore, we cannot predict which ligand initiates signal transduction. Although we were able to quantify changes in activity caused by illumination and phosphorylation via in vitro activity measurements, the molecular mechanism of this regulation remains elusive. While there are multiple regulation and signal transduction mechanisms published concerning LOV domains [28, 29, 71, 72], the combination of a Rec domain linked to a LOV domain has not been described before. It appears intriguing to consider a potential Rec-signal feeding into the A’α helix of LOV, a proven central element of LOV signaling [73]. Eventually this might then affect the LOV-GGDEF linker in ways similar to LOV-, phytochrome-or Rec-GGDEF couples [31, 32, 74]. To address some of the open questions, we are currently following up on an integrative structural biology approach combining data from X-ray crystallography, *in crystallo* optical spectroscopy and hydrogen deuterium exchange coupled to mass spectrometry analyses to probe in solution conformational dynamics across multiple functional states.

In conclusion, *La*RldC was identified as a particularly interesting dual-sensor diguanylate cyclase, showing activity modulation by both phosphorylation and illumination. Based on these results, the protein can be described as operating with a molecular OR logic. Additional factors, such as pH variations and redox inputs, which are certainly encountered in an in vivo context, can also affect activity, rendering this system a peculiar, multilayered enzyme evolved by nature.

## Methods

### Gene selection and cloning

InterPro and PSI-BLAST were used to identify sequences with a Rec-LOV-GGDEF architecture. Multiple sequence alignments were filtered using conserved residues and motifs of the domains as constraints. We classified groups based on the linker between the LOV and GGDEF domains. Similar to previously characterised LOV-GGDEF homologs [28], we defined three classes with different linker lengths, differing by seven or fourteen residues.

Genes were ordered from Thermo Fisher Scientific and sequences codon-optimised for expression in *E. coli*. They were cloned into a pET-M11 vector with an N-terminal His-tag and a TEV protease cleavage site using the NEBuilder HiFi DNA Assembly Cloning Kit to assemble the plasmids.

### In vivo screening

Plasmids encoding the RldC homologs were transformed into *E. coli* BL21 (DE3) and plated on LB Kan plates. After overnight incubation at 37 °C, multiple colonies were used to inoculate a 3 mL liquid culture. The culture was grown at 37 °C to an OD600 of at least 0.5. A volume of 0.5 mL with an OD600 of 0.5 was centrifuged briefly to pellet the cells. The supernatant was discarded, and the cells were resuspended in 25 µL LB medium. Under non-actinic conditions, 2 µL of the resuspended cells were spotted on a Congo Red screening plate (LB (Luria/Miller); 60 mg/L Congo red; 25 mg/L Coomassie G-250; 20 g/L Agar). Plates were wrapped in thin foil and kept dark or illuminated using an LED light panel (Lunartec). The screening plates were incubated at 20 °C for at least 12 hours. Afterwards, plates were kept at 4 °C until pictures were taken with a Canon EOS 1100D, typically after 3-4 days.

### Protein production and purification

To produce the various RldC homologs, we transformed the plasmids described above into *E. coli* BL21 (DE3). The preculture was grown to an OD600 of 1. The main cultures were inoculated with an OD600 of 0.02 and subsequently grown to an OD600 of 0.5-0.6. Afterwards, the cultures were transferred to 16 °C, and after 45 minutes, IPTG was added to a final concentration of 0.1 mM. Cells were harvested after approximately 18 hours (15 min, 4 °C, 3850 × g) and stored at -20 °C until purification.

Cell pellets were resuspended in 5 mL lysis buffer per gram wet cell weight (*Ap, Cp* and *La*: 50 mM HEPES pH 7.0, 500 mM NaCl, 2 mM MgCl2, 30 mM Imidazole; *Ls* and *Ro*: 50 mM CHES pH 9.0, 300 mM NaCl, 2 mM MgCl2, 30 mM Imidazole). Lysozyme (0.5 mg/mL), Protease Inhibitor Cocktail (Roche cOmplete^™^, EDTA free) and DNase I (Roche, grade II) were added to assist cell disruption. After 40-60 minutes of resuspending at room temperature, the cells were disrupted using a B. Braun Labsonic U sonicator (3 × 5 minutes). The suspension was centrifuged (45 min, 4 °C, 37,000 × g) to obtain the supernatant. The supernatant was carefully removed and loaded onto a gravity flow Ni-NTA column (Macherey-Nagel Protino, 35 mL, packed with Cytiva Ni Sepharose 6 Fast). 3 CVs of lysis buffer supplemented with 100 µM FMN (Panreac, AppliChem) were used to ensure even cofactor loading. Then, 6 CVs of lysis buffer containing 40 mM imidazole were used as wash buffer. The protein was eluted using lysis buffer with 250 mM imidazole. Overnight dialysis, lowering the Imidazole concentration to 45 mM, was coupled with tag removal using TEV protease (ratio 1:15 TEV:target protein). The protein solution was then reloaded onto a Ni-NTA column to remove the His-tagged TEV protease and the cleaved tag. The flow-through was concentrated using an Amicon (Merck, 30 kDa MWCO, 3000 × g, 4 °C), followed by size exclusion chromatography as a polishing step (Cytiva, S200 increase 10/300 GL).

To remove bound c-di-GMP, the phosphodiesterase RocR, purified according to Yang et al. [41], was used. Estimating the protein-to-c-di-GMP ratio to be around 1:1, we mixed RocR and the RldC homolog at a 1:5 ratio. To increase turnover, the reaction buffer was supplemented with 100 mM MgCl2. Complete removal was confirmed via HPLC analysis of a denatured reaction mixture sample (conditions similar to the kinetic measurements described below). Afterwards, RocR was separated via affinity Ni-NTA affinity chromatography. For *La*RldC, the addition of 1 mM TCEP during and after c-di-GMP removal was necessary to prevent intermolecular disulfide formation.

### Native mass measurements

The native MS analysis method used in our lab is described in Li et al. (2026) [75]. Briefly, 50 pmol of protein samples (5 µL at 10 µM concentration) were desalted on an AdvanceBio SEC 300 Å (2.7 µm, 4.6 × 50 mm, Agilent) size-exclusion HPLC-column equilibrated in 150 mM ammonium acetate, pH 7. A Shimadzu Nexera UHPLC system was used for automated injections and desalting via the column at a flow rate of 0.2 mL/min. The eluting proteins were directly injected into a Bruker maxis II ultra-high resolution Q-TOF device and the elution volumes containing salts were diverted to the waste valve. Mass spectra were acquired in an m/z range between 500 and 10,000 with 2.5 bar nebuliser pressure and 8 L/min dry gas setting. The source temperature was set to 325 °C and isCID energy to 100 eV. Transfer time and pre-pulse storage were optimised for nMS with 180 µs and 40 µs, respectively. Calibration of the mass spectrometer in the m/z range from 1822 to 5460 was performed using the ESI-tune mix at elevated concentrations, allowing assignments of dimeric and trimeric species. Data was analysed using DataAnalysis (Bruker) and maximum entropy deconvolution of the protein signals.

### Phosphorylation and intact mass measurements

Ammonium hydrogen phosphoramidate was synthesised as previously described [76]. Proteins at 10 µM concentration were incubated with phosphodonor for specified times at 20 °C. Until measurement, the samples were kept at 4°C. Intact mass measurements were performed in-house as previously described by Tran et. al. [46] on a Bruker Impact II QTOF mass spectrometer. Briefly, 5 µl samples were desalted on a Shim-pack Scepter C4-300 (G) column (3 µM) by washing with 1 % ACN in the presence of 0.1 % formic acid. A gradient from 1-95 % ACN was used to elute the proteins. Integration and maximum entropy deconvolution were performed in Bruker “DataAnalysis”.

### Phos-Tag Gels

Phos-Tag™ acrylamide was purchased from FUJIFILM Wako Pure Chemical Corporation and used to prepare Zn2+-Phos-Tag™ SDS-Gels with 6 % acrylamide. Similar to intact mass measurements, proteins were incubated with 50 mM PA at 10 µM protein concentration. After 1 hour of incubation, reactions were stopped by adding 3x sample buffer (according to Wakos Phos-Tag™ Guidebook, but without reducing agent). Samples were loaded without the usual heating denaturation and run at 20°C with constant voltage (100 V).

### Spectroscopic measurements

Spectroscopic measurements were performed using a Specord 200 PLUS dual-beam UV-Vis spectrophotometer (Analytik Jena). Proteins were diluted to 5 µM with storage buffer (Table S.2). The first dark spectrum was recorded under non-actinic conditions. To shift proteins into the light state, a ThorLabs 455 nm or 470 nm mounted LED (3 mW/cm^2^) was used. Samples were illuminated for specified times, after which spectra were measured. Spectra were recorded until no further light induced change was observed. Dark-state recovery was determined by collecting spectra from 300 to 650 nm at predetermined time intervals to detect potential aggregation effects during long measurement times.

### Kinetic measurements

Protein stocks were diluted with reaction buffer (100 mM Tris pH 8.0, 500 mM NaCl, 10 mM MgCl_2_, 1 mM TCEP) to 2 µM. Dark state samples were started by the addition of GTP. For the individual time points, 40 µL of the reaction was withdrawn and stopped by the addition of EDTA and additional heat denaturation (96 °C, 1 min). For light state measurements, samples were pre-illuminated for time intervals based on the initial spectroscopic characterisation (LaRldC: 5 min; *Cp*RldC: 90 sec; *Ro*RldC: 90 sec), before adding GTP to start the reaction. Samples were centrifuged for 10 min at 13,000 rpm before HPLC analysis. The samples were separated isocratically (10 mM K_2_HPO_4_; 1 mM EDTA) on a C18 column (Bischoff ProntoSIL, 1250-5-C18 ace-EPS, 100 × 4.6 mm) at 25°C. Nucleotides were detected based on 254 nm absorbance, with quantification based on relative peak heights. All measurements were performed in triplicate to evaluate the standard deviation.

## Acknowledgements

We thank Roland C. Fischer for the synthesis of ammonium hydrogen phosphoramidate. We appreciate the critical review of the manuscript by Peter Macheroux. We also thank Philipp Pelzmann for performing intact and native MS measurements.

## Funding and additional information

M. F. acknowledges support as an associate student from the Austrian Science Fund (FWF) grant DOC-130. This research was funded in whole or in part by the Austrian Science Fund (FWF) [grant doi: 10.55776/P34387; to A. W.].

## Notes

### Competing Interest Statement

The authors have declared no competing interest.

## References

[1] Nicolas Papon, Ann M. Stock, “Two-component systems”, Current biology CB, Vol. 29, No. 15, R724–R725.

[2] Haike Antelmann, John D. Helmann, “Thiol-based redox switches and gene regulation”, Antioxidants & redox signaling, Vol. 14, No. 6, pp. 1049–1063, 2011.

[3] Daniel G. Isom, Vishwajith Sridharan, Rachael Baker, Sarah T. Clement, David M. Smalley, Henrik G. Dohlman, “Protons as second messenger regulators of G protein signaling”, Molecular cell, Vol. 51, No. 4, pp. 531–538.

[4] Mark Gomelsky, Gabriele Klug, “BLUF: a novel FAD-binding domain involved in sensory transduction in microorganisms”, Trends in biochemical sciences, Vol. 27, No. 10, pp. 497–500, 2002.

[5] Sean Crosson, Keith Moffat, “Photoexcited structure of a plant photoreceptor domain reveals a light-driven molecular switch”, The Plant cell, Vol. 14, No. 5, pp. 1067–1075, 2002.

[6] Jason A. Papin, Tony Hunter, Bernhard O. Palsson, Shankar Subramaniam, “Reconstruction of cellular signalling networks and analysis of their properties”, Nature reviews. Molecular cell biology, Vol. 6, No. 2, pp. 99–111.

[7] Roby P. Bhattacharyya, Attila Reményi, Brian J. Yeh, Wendell A. Lim, “Domains, motifs, and scaffolds: the role of modular interactions in the evolution and wiring of cell signaling circuits”, Annual review of biochemistry, Vol. 75, pp. 655–680, 2006.

[8] Vanessa I. Francis, Steven L. Porter, “Multikinase Networks: Two-Component Signaling Networks Integrating Multiple Stimuli”, Annual review of microbiology, Vol. 73, pp. 199–223, 2019.

[9] Thomas Guest, James R. J. Haycocks, Gemma Z. L. Warren, David C. Grainger, “Genome-wide mapping of Vibrio cholerae VpsT binding identifies a mechanism for c-di-GMP homeostasis”, Nucleic acids research, Vol. 50, No. 1, pp. 149–159, 2022.

[10] Liang Nie, Yujie Xiao, Tiantian Zhou, Haoqi Feng, Meina He, Qingyuan Liang, Kexin Mu, Hailing Nie, Qiaoyun Huang, Wenli Chen, “Cyclic di-GMP inhibits nitrate assimilation by impairing the antitermination function of NasT in Pseudomonas putida”, Nucleic acids research, Vol. 52, No. 1, pp. 186–203, 2024.

[11] Ning-Ning Liu, Meng-Lin Li, Wen-Tao Shi, Jian Jiao, Yan-Hui Xu, Yu Tian, Jia-Ning Guo, Yu-Qing Chen, Huan Tong, Chang-Fu Tian, “Cyclic-di-GMP interferes with DNA-MucR-DNA bridging to derepress genes targeted by the xenogeneic silencer MucR”, Nucleic acids research, Vol. 53, No. 20, 2025.

[12] Regine Hengge, Angelika Gründling, Urs Jenal, Robert Ryan, Fitnat Yildiz, “Bacterial Signal Transduction by Cyclic Di-GMP and Other Nucleotide Second Messengers”, Journal of bacteriology, Vol. 198, No. 1, pp. 15–26, 2016.

[13] Rita Tamayo, Jason T. Pratt, Andrew Camilli, “Roles of cyclic diguanylate in the regulation of bacterial pathogenesis”, Annual review of microbiology, Vol. 61, pp. 131–148, 2007.

[14] Nancy Obeng, Anna Czerwinski, Daniel Schütz, Jan Michels, Jan Leipert, Florence Bansept, María J. García García, Thekla Schultheiß, Melinda Kemlein, Janina Fuß, Andreas Tholey, Arne Traulsen, Holger Sondermann, Hinrich Schulenburg, “Bacterial c-di-GMP has a key role in establishing host-microbe symbiosis”, Nature microbiology, Vol. 8, No. 10, pp. 1809–1819.

[15] Gen Enomoto, Thomas Wallner, Annegret Wilde, “Control of light-dependent behaviour in cyanobacteria by the second messenger cyclic di-GMP”, microLife, Vol. 4, uqad019, 2023.

[16] Holger Sondermann, Nicholas J. Shikuma, Fitnat H. Yildiz, “You’ve come a long way: c-di-GMP signaling”, Current opinion in microbiology, Vol. 15, No. 2, pp. 140–146, 2012.

[17] Ralf Paul, Stefan Weiser, Nicholas C. Amiot, Carmen Chan, Tilman Schirmer, Bernd Giese, Urs Jenal, “Cell cycle-dependent dynamic localization of a bacterial response regulator with a novel di-guanylate cyclase output domain”, Genes & development, Vol. 18, No. 6, pp. 715–727, 2004.

[18] David G. Welkie, Benjamin E. Rubin, Spencer Diamond, Rachel D. Hood, David F. Savage, Susan S. Golden, “A Hard Day’s Night: Cyanobacteria in Diel Cycles”, Trends in microbiology, Vol. 27, No. 3, pp. 231–242, 2019.

[19] Jia Guo, Amelia Y. Nguyen, Ziyu Dai, Dian Su, Matthew J. Gaffrey, Ronald J. Moore, Jon M. Jacobs, Matthew E. Monroe, Richard D. Smith, David W. Koppenaal, Himadri B. Pakrasi, Wei-Jun Qian, “Proteome-wide light/dark modulation of thiol oxidation in cyanobacteria revealed by quantitative site-specific redox proteomics”, Molecular & cellular proteomics MCP, Vol. 13, No. 12, pp. 3270–3285.

[20] Niall M. Mangan, Avi Flamholz, Rachel D. Hood, Ron Milo, David F. Savage, “pH determines the energetic efficiency of the cyanobacterial CO2 concentrating mechanism”, Proceedings of the National Academy of Sciences of the United States of America, Vol. 113, No. 36, E5354-62.

[21] Gen Enomoto, Ni-Ni-Win, Rei Narikawa, Masahiko Ikeuchi, “Three cyanobacteriochromes work together to form a light color-sensitive input system for c-di-GMP signaling of cell aggregation”, Proceedings of the National Academy of Sciences of the United States of America, Vol. 112, No. 26, pp. 8082–8087, 2015.

[22] Philipp Savakis, Sven de Causmaecker, Veronika Angerer, Ulrike Ruppert, Katrin Anders, Lars-Oliver Essen, Annegret Wilde, “Light-induced alteration of c-di-GMP level controls motility of Synechocystis sp. PCC 6803”, Molecular microbiology, Vol. 85, No. 2, pp. 239–251, 2012.

[23] Marco Agostoni, Benjamin J. Koestler, Christopher M. Waters, Barry L. Williams, Beronda L. Montgomery, “Occurrence of cyclic di-GMP-modulating output domains in cyanobacteria: an illuminating perspective”, mBio, Vol. 4, No. 4.

[24] Rong Gao, Sophie Bouillet, Ann M. Stock, “Structural Basis of Response Regulator Function”, Annual review of microbiology, Vol. 73, pp. 175–197, 2019.

[25] Yingpeng Xie, Jingwei Li, Yiqing Ding, Xiaolong Shao, Yue Sun, Fangzhou Xie, Shiyi Liu, Shaojun Tang, Xin Deng, “An atlas of bacterial two-component systems reveals function and plasticity in signal transduction”, Cell reports, Vol. 41, No. 3, p. 111502, 2022.

[26] T. E. Swartz, S. B. Corchnoy, J. M. Christie, J. W. Lewis, I. Szundi, W. R. Briggs, R. A. Bogomolni, “The photocycle of a flavin-binding domain of the blue light photoreceptor phototropin”, Journal of Biological Chemistry, Vol. 276, No. 39, pp. 36493–36500, 2001.

[27] Spencer T. Glantz, Eric J. Carpenter, Michael Melkonian, Kevin H. Gardner, Edward S. Boyden, Gane K.-S. Wong, Brian Y. Chow, “Functional and topological diversity of LOV domain photoreceptors”, Proceedings of the National Academy of Sciences of the United States of America, Vol. 113, No. 11, E1442-51, 2016.

[28] Uršula Vide, Dženita Kasapović, Maximilian Fuchs, Martin P. Heimböck, Massimo G. Totaro, Elfriede Zenzmaier, Andreas Winkler, “Illuminating the inner workings of a natural protein switch: Blue-light sensing in LOV-activated diguanylate cyclases”, Science advances, Vol. 9, No. 31, eadh4721, 2023.

[29] Danielle Swingle, Leah Epstein, Ramisha Aymon, Eta A. Isiorho, Rinat R. Abzalimov, Denize C. Favaro, Kevin H. Gardner, “Variations in kinase and effector signaling logic in a bacterial two component signaling network”, The Journal of biological chemistry, Vol. 301, No. 6, p. 108534.

[30] Josiah P. Zayner, Chloe Antoniou, Tobin R. Sosnick, “The amino-terminal helix modulates light-activated conformational changes in AsLOV2”, Journal of molecular biology, Vol. 419, 1-2, pp. 61– 74, 2012.

[31] Uršula Vide, Gabriela Shickle, Julia Schwekendiek, Andreas Winkler, “Coiled-coil register transitions and coupling with the effector’s inhibitory site enables high fold changes in blue light-regulated diguanylate cyclases”, The Journal of biological chemistry, Vol. 302, No. 1, p. 111020.

[32] Raphael D. Teixeira, Fabian Holzschuh, Tilman Schirmer, “Activation mechanism of a small prototypic Rec-GGDEF diguanylate cyclase”, Nature communications, Vol. 12, No. 1, p. 2162, 2021.

[33] Geoffrey Gourinchas, Stefan Etzl, Christoph Göbl, Uršula Vide, Tobias Madl, Andreas Winkler, “Long-range allosteric signaling in red light-regulated diguanylyl cyclases”, Science advances, Vol. 3, No. 3, e1602498, 2017.

[34] Jaynee E. Hart, Kevin H. Gardner, “Lighting the way: Recent insights into the structure and regulation of phototropin blue light receptors”, The Journal of biological chemistry, Vol. 296, p. 100594, 2021.

[35] Ophilia I. L. Mawphlang, M. Bharatheeswaran, Lingaraj Sahoo, Highland Kayang, Eros Kharshiing, “Cloning and expression analyses of the twin LOV protein (LLP) gene under varying light, water-deficit and humidity conditions in the small-fruited tomato (Solanum lycopersicum var. cerasiforme)”, Plant Gene, Vol. 45, p. 100560, 2026.

[36] Robert B. Bourret, “Receiver domain structure and function in response regulator proteins”, Current opinion in microbiology, Vol. 13, No. 2, pp. 142–149, 2010.

[37] Linda Truebestein, Thomas A. Leonard, “Coiled-coils: The long and short of it”, BioEssays news and reviews in molecular, cellular and developmental biology, Vol. 38, No. 9, pp. 903–916, 2016.

[38] Efrosini Moutevelis, Derek N. Woolfson, “A periodic table of coiled-coil protein structures”, Journal of molecular biology, Vol. 385, No. 3, pp. 726–732.

[39] Roger S. Annika Cimdins, “Semiquantitative Analysis of the Red, Dry, and Rough Colony Morphology of Salmonella enterica Serovar Typhimurium and Escherichia coli Using Congo Red”, Vol. 2017.

[40] Beat Christen, Matthias Christen, Ralf Paul, Franziska Schmid, Marc Folcher, Paul Jenoe, Markus Meuwly, Urs Jenal, “Allosteric Control of Cyclic di-GMP Signaling”, Journal of Biological Chemistry, Vol. 281, No. 42, pp. 32015–32024.

[41] Feng Rao, Ye Yang, Yaning Qi, Zhao-Xun Liang, “Catalytic mechanism of cyclic di-GMP-specific phosphodiesterase: a study of the EAL domain-containing RocR from Pseudomonas aeruginosa”, Journal of bacteriology, Vol. 190, No. 10, pp. 3622–3631, 2008.

[42] Josiah P. Zayner, Tobin R. Sosnick, “Factors that control the chemistry of the LOV domain photocycle”, PloS one, Vol. 9, No. 1, e87074, 2014.

[43] Raoul E. Herzog, Isabelle F. Harvey-Seutcheu, Philipp Janke, Wenzhao Dai, Paul M. Fischer, Peter Hamm, Philipp J. Heckmeier, “Evolution and design shape protein dynamics in LOV domains - spanning picoseconds to days”, Journal of molecular biology, Vol. 438, No. 5, p. 169599, 2025.

[44] T. L. Mayover, C. J. Halkides, R. C. Stewart, “Kinetic characterization of CheY phosphorylation reactions: comparison of P-CheA and small-molecule phosphodonors”, Biochemistry, Vol. 38, No. 8, pp. 2259–2271, 1999.

[45] R. B. Bourret, S. K. Drake, S. A. Chervitz, M. I. Simon, J. J. Falke, “Activation of the phosphosignaling protein CheY. II. Analysis of activated mutants by 19F NMR and protein engineering”, Journal of Biological Chemistry, Vol. 268, No. 18, pp. 13089–13096.

[46] Quang H. Tran, Oliver M. Eder, Andreas Winkler, “Dynamics-driven allosteric stimulation of diguanylate cyclase activity in a red light-regulated phytochrome”, The Journal of biological chemistry, Vol. 300, No. 5, p. 107217.

[47] Cornelia Böhm, Nikolina Todorović, Marco Balasso, Geoffrey Gourinchas, Andreas Winkler, “The PHY Domain Dimer Interface of Bacteriophytochromes Mediates Cross-talk between Photosensory Modules and Output Domains”, Journal of molecular biology, Vol. 433, No. 15, p. 167092.

[48] Heewhan Shin, Zhong Ren, Xiaoli Zeng, Sepalika Bandara, Xiaojing Yang, “Structural basis of molecular logic OR in a dual-sensor histidine kinase”, Proceedings of the National Academy of Sciences of the United States of America, Vol. 116, No. 40, pp. 19973–19982, 2019.

[49] Ayumu Kobayashi, Masamune Nakamura, Masaru Tsujii, Kohei Makino, Tatsuya Nagayama, Kensuke Nakamura, Kei Nanatani, Kera Kota, Yuki Furuuchi, Shunsuke Kayamori, Tadaomi Furuta, Iwane Suzuki, Yoshihiro Hayakawa, Ellen Tanudjaja, Yasuhiro Ishimaru, Nobuyuki Uozumi, “Two cyanobacterial response regulators with diguanylate cyclase activity, Rre2 and Rre8, participate in biofilm formation”, Molecular microbiology, Vol. 119, No. 5, pp. 599–611, 2023.

[50] Huan-Xiang Zhou, Xiaodong Pang, “Electrostatic Interactions in Protein Structure, Folding, Binding, and Condensation”, Chemical reviews, Vol. 118, No. 4, pp. 1691–1741, 2018.

[51] Claudia M. Cremers, Ursula Jakob, “Oxidant sensing by reversible disulfide bond formation”, The Journal of biological chemistry, Vol. 288, No. 37, pp. 26489–26496.

[52] Marcele P. Martins, Gustavo H. Martins, Felipe J. Fuzita, João P. M. Spadeto, Renan Y. Miyamoto, Felippe M. Colombari, Fabiane Stoffel, Luciano G. Dolce, Camila R. D. Santos, Rodrigo S. A. Streit, Antônio C. Borges, Rafael H. Galinari, Yoshihisa Yoshimi, Paul Dupree, Gabriela F. Persinoti, Mariana A. B. Morais, Mario T. Murakami, “A disulfide redox switch mechanism regulates glycoside hydrolase function”, Nature communications, Vol. 17, No. 1, p. 45.

[53] Bin Li, Minshik Jo, Jianxin Liu, Jiayi Tian, Robert Canfield, Jennifer Bridwell-Rabb, “Structural and mechanistic basis for redox sensing by the cyanobacterial transcription regulator RexT”, Communications biology, Vol. 5, No. 1, p. 275.

[54] Aba Losi, Wolfgang Gärtner, “Solving Blue Light Riddles: New Lessons from Flavin-binding LOV Photoreceptors”, Photochemistry and photobiology, Vol. 93, No. 1, pp. 141–158.

[55] Brian D. Zoltowski, Takato Imaizumi, “Structure and Function of the ZTL/FKF1/LKP2 Group Proteins in Arabidopsis”, The Enzymes, Vol. 35, pp. 213–239, 2014.

[56] Ashutosh Pudasaini, Brian D. Zoltowski, “Zeitlupe senses blue-light fluence to mediate circadian timing in Arabidopsis thaliana”, Biochemistry, Vol. 52, No. 40, pp. 7150–7158, 2013.

[57] Ashutosh Pudasaini, Jae S. Shim, Young H. Song, Hua Shi, Takatoshi Kiba, David E. Somers, Takato Imaizumi, Brian D. Zoltowski, “Kinetics of the LOV domain of ZEITLUPE determine its circadian function in Arabidopsis”, eLife, Vol. 6, 2017.

[58] Julia Dietler, Renate Gelfert, Jennifer Kaiser, Veniamin Borin, Christian Renzl, Sebastian Pilsl, Américo T. Ranzani, Andrés García de Fuentes, Tobias Gleichmann, Ralph P. Diensthuber, Michael Weyand, Günter Mayer, Igor Schapiro, Andreas Möglich, “Signal transduction in light-oxygen-voltage receptors lacking the active-site glutamine”, Nature communications, Vol. 13, No. 1, p. 2618, 2022.

[59] Salomon, M., Christie, J. M., Knieb, E., Lempert, U. & Briggs, W. R., “Photochemical and Mutational Analysis of the FMN-Binding Domains of the Plant Blue Light Receptor”, Biochemistry, Vol. 2000, No. 39, pp. 9401–9410, 2000.

[60] Masahiro Kasahara, Mayumi Torii, Akimitsu Fujita, Kengo Tainaka, “FMN binding and photochemical properties of plant putative photoreceptors containing two LOV domains, LOV/LOV proteins”, The Journal of biological chemistry, Vol. 285, No. 45, pp. 34765–34772, 2010.

[61] Dasvit Shetty, Linda J. Kenney, “A pH-sensitive switch activates virulence in Salmonella”, eLife, Vol. 12.

[62] Igor Dikiy, Uthama R. Edupuganti, Rinat R. Abzalimov, Peter P. Borbat, Madhur Srivastava, Jack H. Freed, Kevin H. Gardner, “Insights into histidine kinase activation mechanisms from the monomeric blue light sensor EL346”, Proceedings of the National Academy of Sciences of the United States of America, Vol. 116, No. 11, pp. 4963–4972, 2019.

[63] Mohamed Watad, Lukas Korf, Wieland Steinchen, Filipp Bezold, Marian S. Vogt, Po H. Wang, Leon Selbach, Sebastian Hepp, Luis Gayermann, Marleen van Wolferen, Xing Ye, Sonja-Verena Albers, Lars-Oliver Essen, “Phosphorylation-driven conformational switching of the ArnA-ArnB complex involved in archaeal motility regulation”, Frontiers in microbiology, Vol. 16, p. 1717585, 2025.

[64] Udo Heintz, Ilme Schlichting, “Blue light-induced LOV domain dimerization enhances the affinity of Aureochrome 1a for its target DNA sequence”, eLife, Vol. 5, e11860, 2016.

[65] Robert Lindner, Elisabeth Hartmann, Miroslaw Tarnawski, Andreas Winkler, Daniel Frey, Jochen Reinstein, Anton Meinhart, Ilme Schlichting, “Photoactivation Mechanism of a Bacterial Light-Regulated Adenylyl Cyclase”, Journal of molecular biology, Vol. 429, No. 9, pp. 1336–1351, 2017.

[66] Andreas Winkler, Udo Heintz, Robert Lindner, Jochen Reinstein, Robert L. Shoeman, Ilme Schlichting, “A ternary AppA-PpsR-DNA complex mediates light regulation of photosynthesis-related gene expression”, Nature structural & molecular biology, Vol. 20, No. 7, pp. 859–867, 2013.

[67] Andreas Winkler, Anikó Udvarhelyi Elisabeth Hartmann, Jochen Reinstein, Andreas Menzel, Robert L. Shoeman, Ilme Schlichting, “Characterization of elements involved in allosteric light regulation of phosphodiesterase activity by comparison of different functional BlrP1 states”, Journal of molecular biology, Vol. 426, No. 4, pp. 853–868, 2014.

[68] M. Jiang, R. B. Bourret, M. I. Simon, K. Volz, “Uncoupled phosphorylation and activation in bacterial chemotaxis. The 2.3 A structure of an aspartate to lysine mutant at position 13 of CheY”, The Journal of biological chemistry, Vol. 272, No. 18, pp. 11850–11855.

[69] Amit A. Upadhyay, Aaron D. Fleetwood, Ogun Adebali, Robert D. Finn, Igor B. Zhulin, “Cache Domains That are Homologous to, but Different from PAS Domains Comprise the Largest Superfamily of Extracellular Sensors in Prokaryotes”, PLoS computational biology, Vol. 12, No. 4, e1004862.

[70] Miguel A. Matilla, Félix Velando, David Martín-Mora, Elizabet Monteagudo-Cascales, Tino Krell, “A catalogue of signal molecules that interact with sensor kinases, chemoreceptors and transcriptional regulators”, FEMS microbiology reviews, Vol. 46, No. 1.

[71] Aditya S. Chaudhari, Adrien Favier, Zahra A. Tehrani, Tomáš Kovaľ, Inger Andersson, Bohdan Schneider, Jan Dohnálek, Jiří Černý, Bernhard Brutscher, Gustavo Fuertes, “Light-dependent flavin redox and adduct states control the conformation and DNA-binding activity of the transcription factor EL222”, Nucleic acids research, Vol. 53, No. 6, 2025.

[72] Julien Herrou, Sean Crosson, “Function, structure and mechanism of bacterial photosensory LOV proteins”, Nature Reviews Microbiology, Vol. 9, No. 10, pp. 713–723, 2011.

[73] Andreas Möglich, Xiaojing Yang, Rebecca A. Ayers, Keith Moffat, “Structure and function of plant photoreceptors”, Annual review of plant biology, Vol. 61, pp. 21–47, 2010.

[74] Geoffrey Gourinchas, Udo Heintz, Andreas Winkler, “Asymmetric activation mechanism of a homodimeric red light-regulated photoreceptor”, eLife, Vol. 7, 2018.

[75] Tuo Li, Pedro A. Sánchez-Murcia, Bernd Nidetzky, “Structural instability impairs function of the UDP-xylose synthase 1 Ile181Asn variant associated with short-stature genetic syndrome in humans”, FEBS letters, 2026.

[76] F. A. Cotton (Ed.), “Inorganic syntheses: Volume XIII”, McGraw Hill, New York, 1971.

