## Supplemental Material for "Molecular logics in dual sensor regulation of enzyme activity – Phosphorylation OR blue-light activation of cyanobacterial diguanylate cyclases"

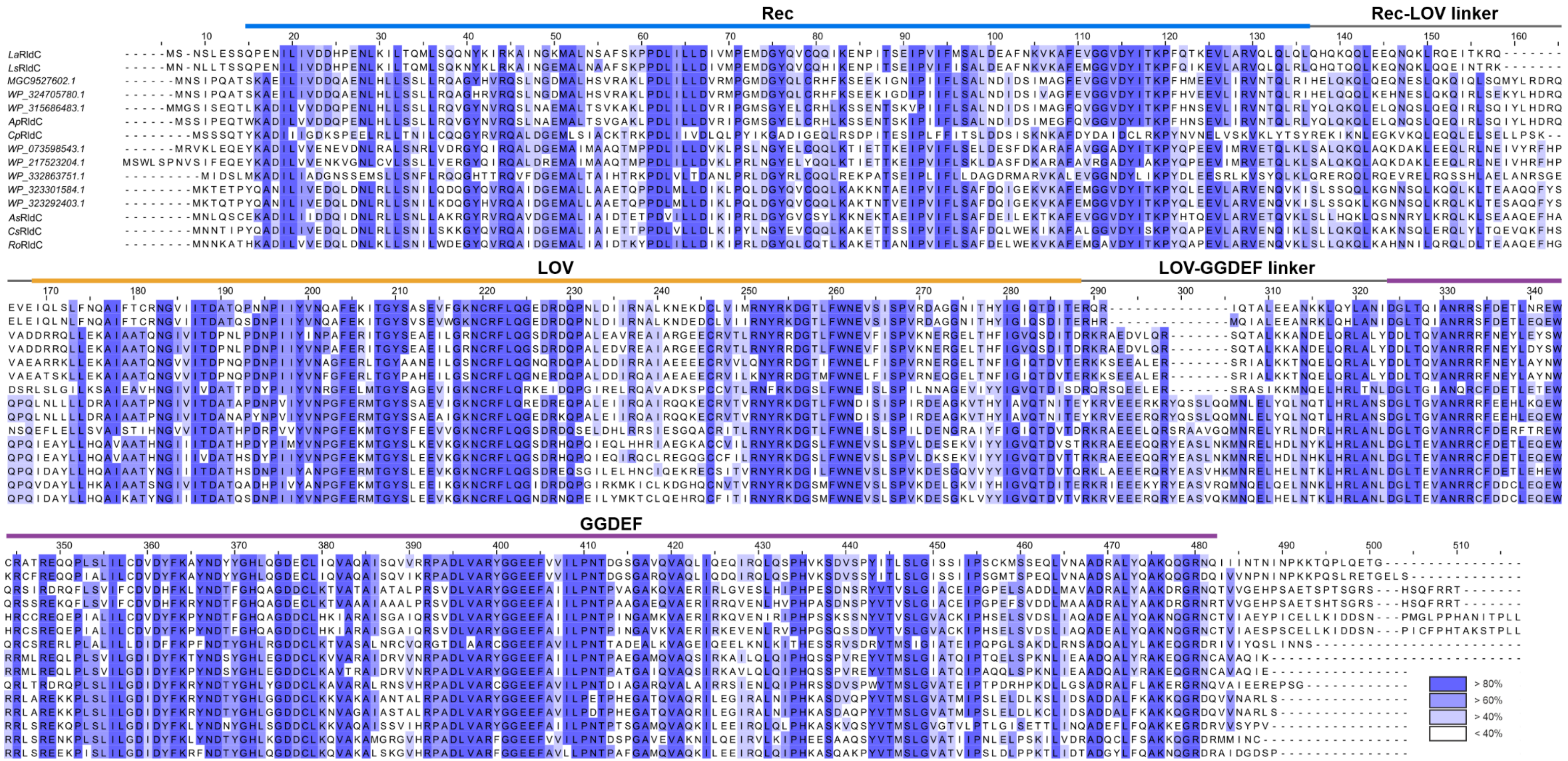


**Supplemental Figure 1: Sequence alignment of selected RldC homologs. The** alignment was filtered for redundancy, eliminating sequences with identities larger than 95%. Individual domains and linker regions are labelled. Colours correspond to to percentage identity as shown in the bottom right.

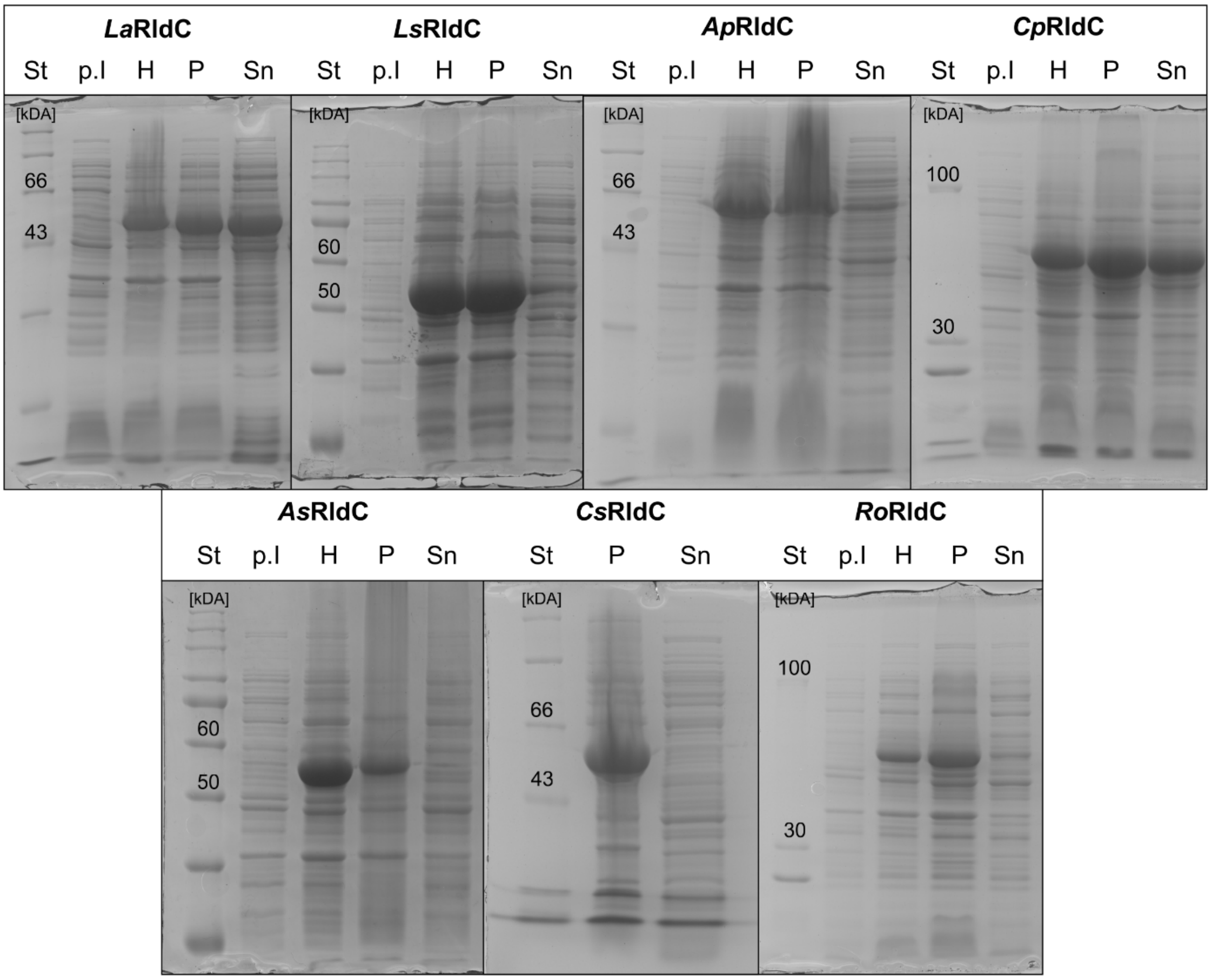


**Supplemental Figure 2: SDS-PAGE analysis of individual homologs.** Gels were used to judge the solubility of individual homologs after overnight protein production. p.I = before induction; H = harvested cells after overnight protein production; P = cell pellet after extraction of soluble fraction; Sn = supernatant after centrifugation containing soluble fractions.

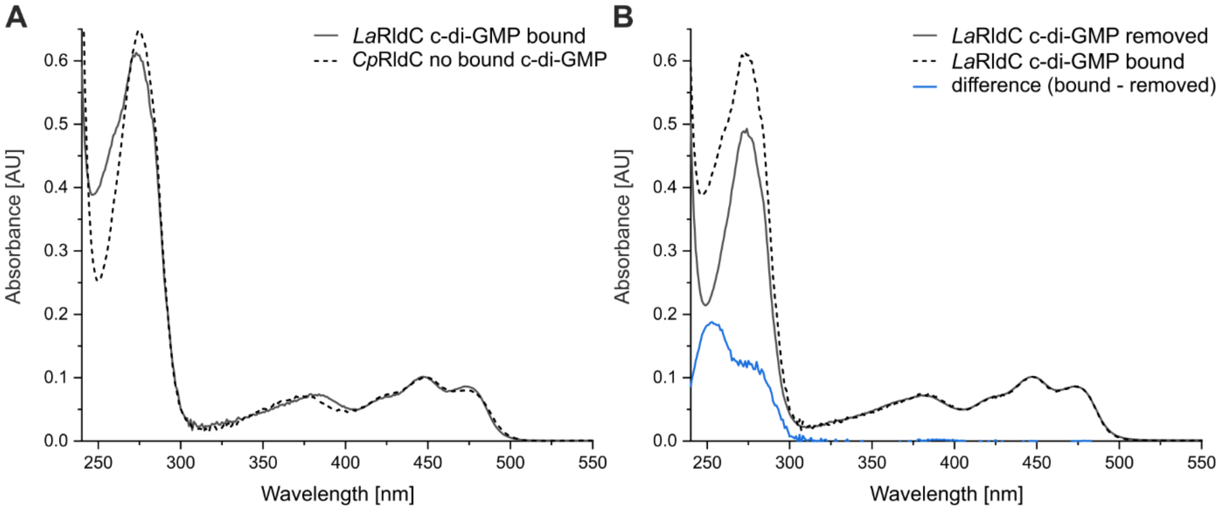


**Supplemental Figure 3: Absorption spectra identifying bound c-di-GMP.** A) UV absorbance differs between *Cp*RldC (no bound c-di-GMP) and *La*RldC (bound c-di-GMP). B) Comparison of *La*RldC before and after removing bound c-di-GMP. The difference spectrum matches the spectrum of guanosine.

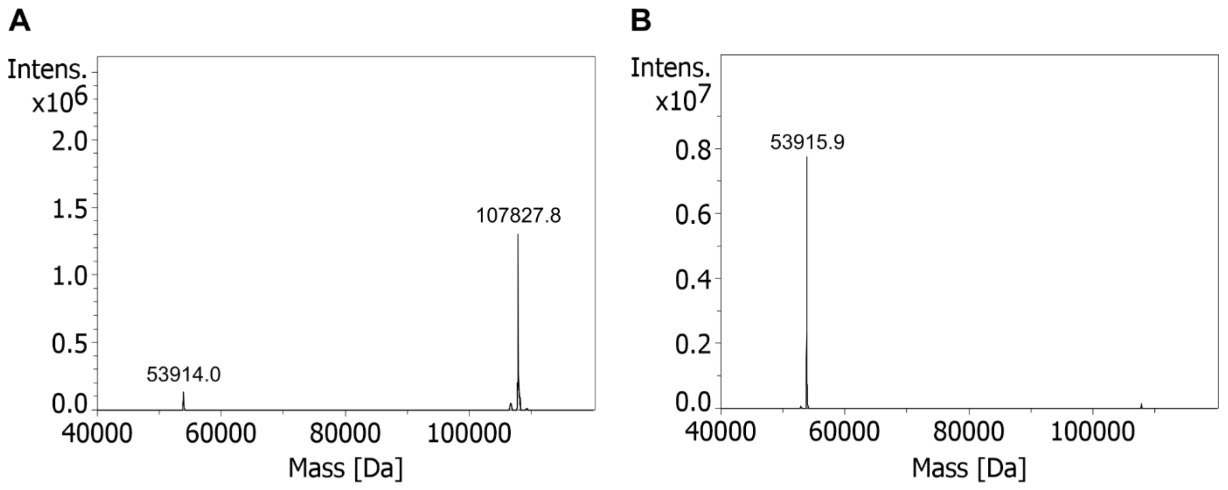


**Supplemental Figure 4: Intact Mass measurements of *La*RldC after removal of the copurified c-di-GMP.** A) Measured after removal of bound c-di-GMP. In addition to the monomer mass, a dimer is also present. The expected mass for the protein is 53,915.8 Da. B) Measurement after the addition of TCEP to the buffer and detection of only the monomer's mass. The minor amount of dimer can be considered an artefact of maximum entropy deconvolution.

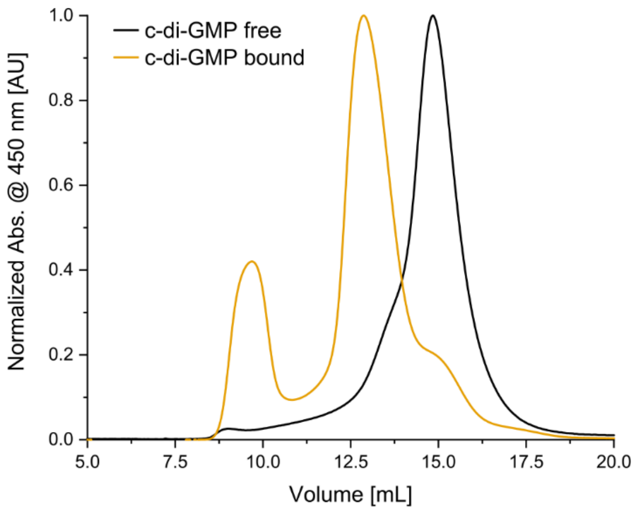


**Supplemental Figure 5: FPLC traces of the *Ro*RldC purification.** The elution volume of the main peak changes once bound c-di-GMP is removed. Purification on a Cytiva S200 increase 10/300 column equilibrated in 10 mM Tris pH 8.0, 500 mM NaCl, 2 mM NaCl.

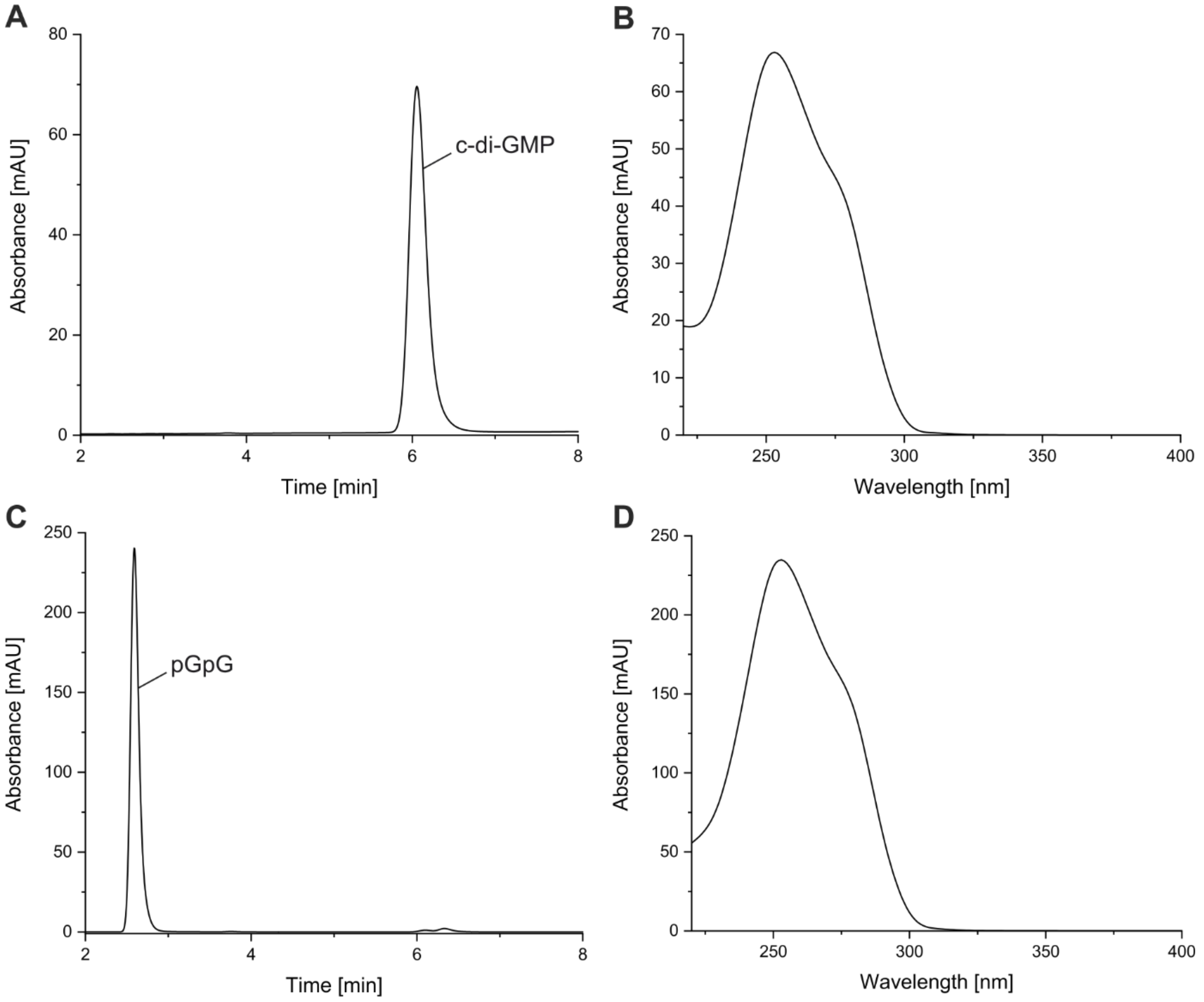


**Supplemental Figure 6: HPLC traces and peak spectra confirming bound or converted c-di-GMP.** A) Denatured sample of *La*RldC showing bound c-di-GMP. B) Absorption spectrum of the peak visible in panel A. C) Denatured sample of *La*RldC after incubation with RocR showing conversion of c-di-GMP to pGpG. D) Absorption spectrum of the pGpG peak shown in C.

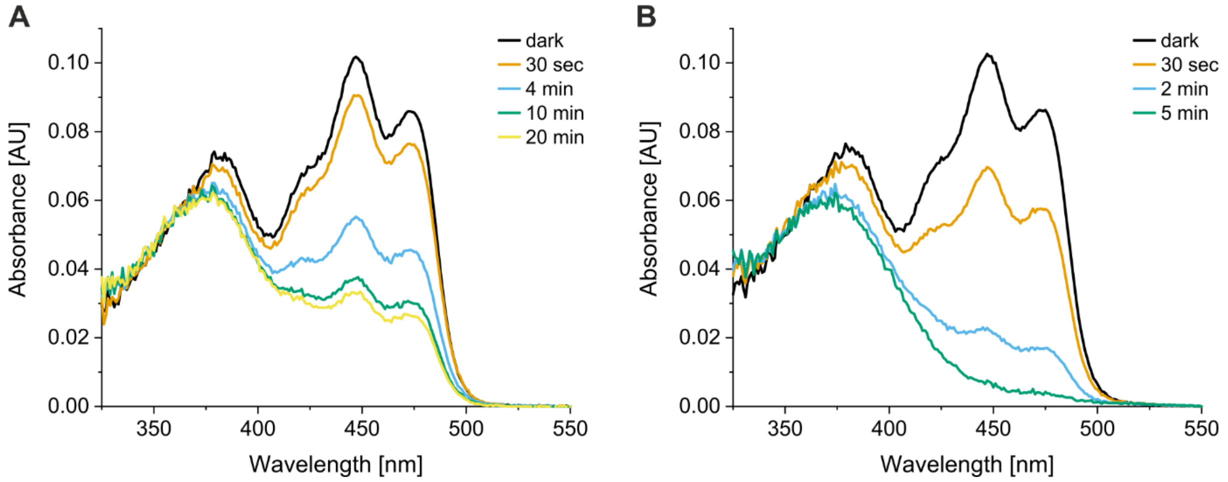


**Supplemental Figure 7: Influence of light quality on dark- to light-state transition of *La*RldC.** A) Sample illuminated with 455 nm blue light LED (ThorLabs, 3 mW/cm$^2$) for indicated amounts of time. B) Illumination with 470 nm blue light LED (ThorLabs, 3 mW/cm$^2$) for specified times.

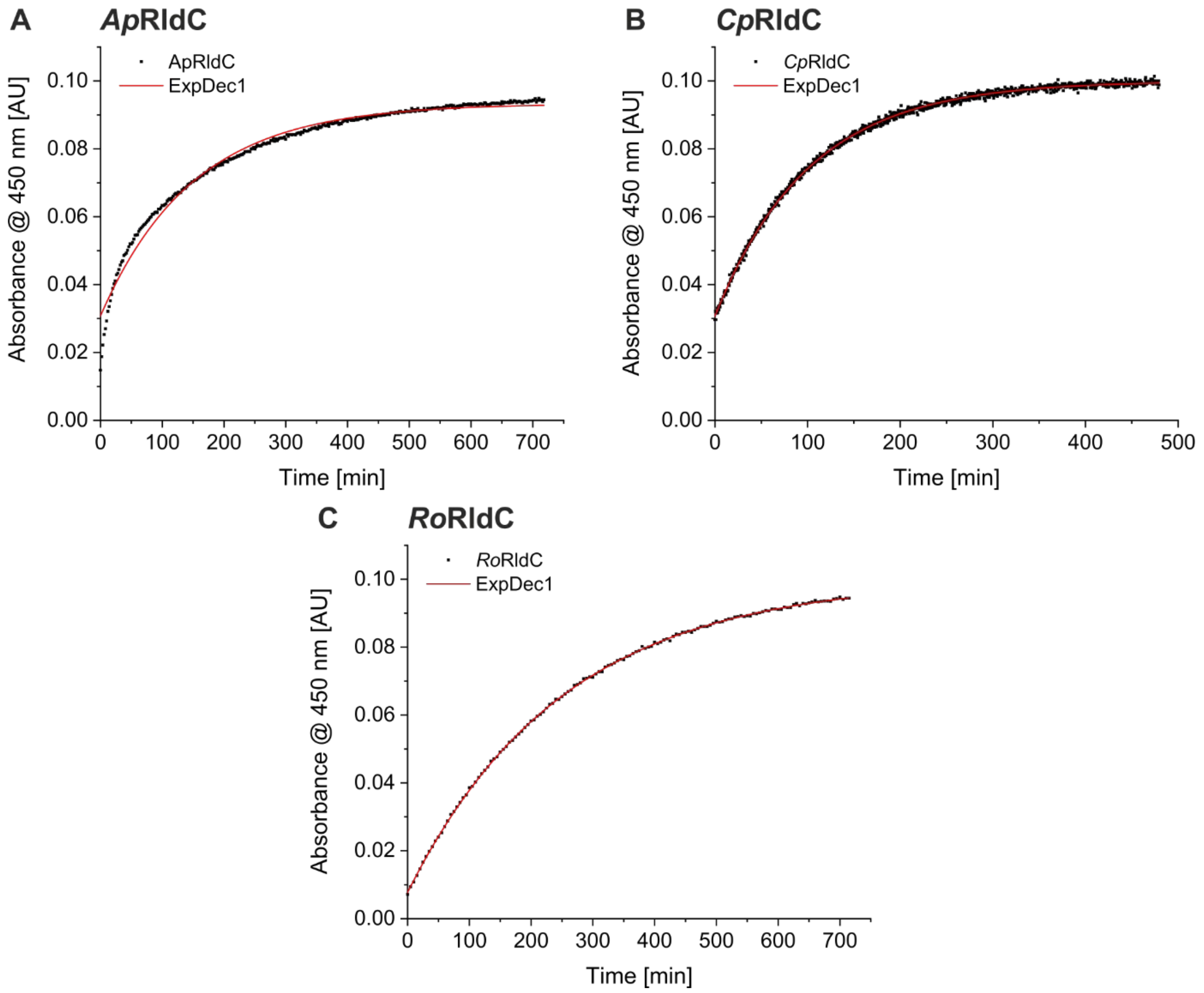


***Supplemental Figure 8:*** **Single exponential fits of thermal recoveries.** A) Shows *Ap*RldC recovery data (black) and exponential fit (red). Equation with rounded parameters $y = -0.06 * e^{(-x/150)} + 0.09$; τ (mean lifetime) = 150 min. B) Recovery data of *Cp*RldC in black with calculated fit in red. $y = -0.10 * e^{(-x/101)} + 0.11$; τ = 101 min. C) Measured recovery data of *Ro*RldC (black) with exponential fit (red). $y = -0.09 * e^{(-x/253)} + 0.1$; τ = 253 min.

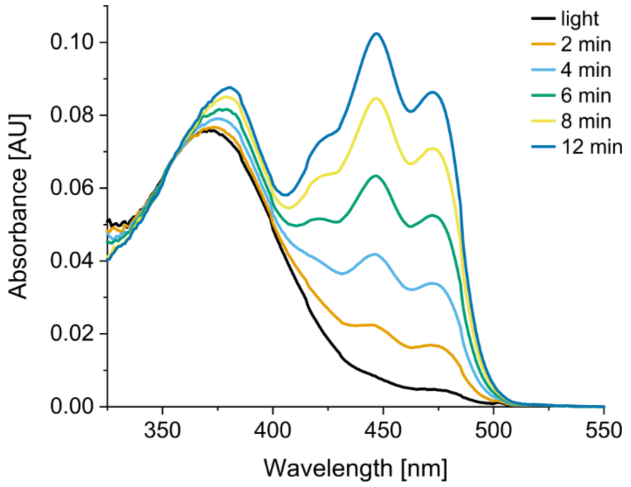


**Supplemental Figure 9: UV illumination of *La*RldC.** Illumination with 395 nm light (2 mW/cm$^2$) switches the protein back to the dark state. Times of illumination are specified in the legend.

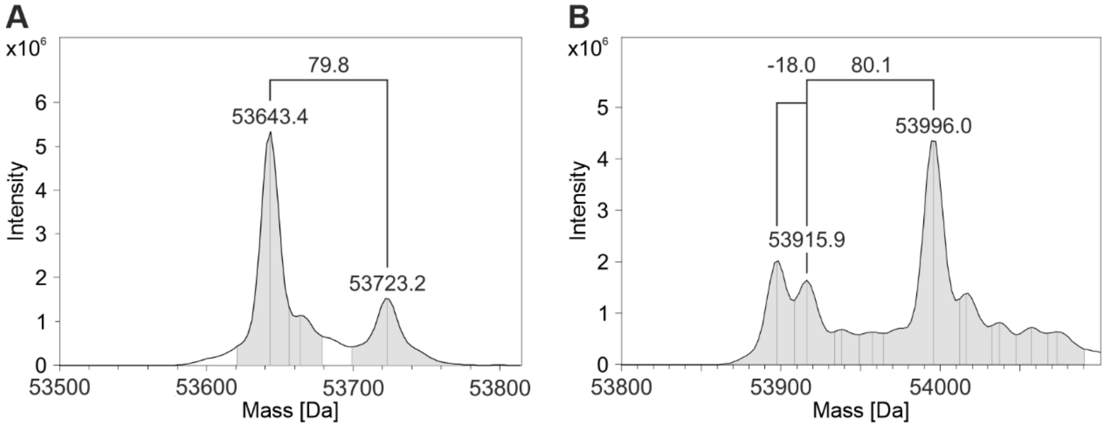


**Supplemental Figure 10: Intact mass measurement after incubation with 50 mM phosphoramidate.** A) *Cp*RldC after multiple hours of incubation at 20 °C. Only moderate amounts of phosphorylated protein were detected. The approximately 80 Da mass difference is characteristic of protein phosphorylation. B) Mass spectrum showing phosphorylation of *La*RldC. The majority of the protein was phosphorylated after 1 h incubation at 20 °C. The peak at -18.0 Da is due to the neutral loss of $H_3PO_4$ from the phosphorylated species, a well-documented phenomenon in phosphorylated proteins [77, 78]

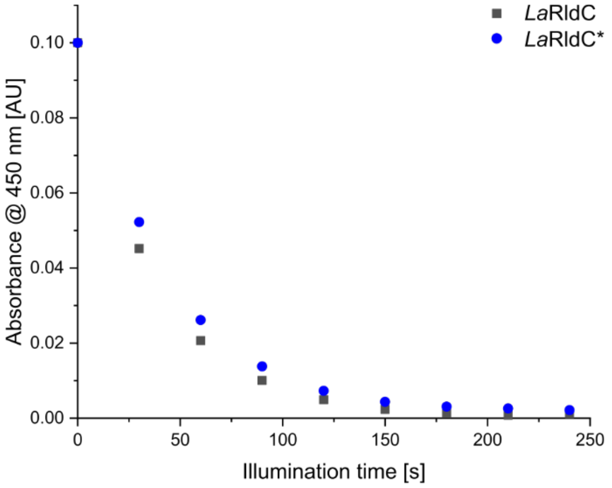


**Supplemental Figure 11: Activation of *La*RldC with and without phosphorylation.** Shows the decrease of the 450 nm absorbance in response to illumination with 470 nm blue light. The phosphorylated sample indicated by * was incubated with 50 mM phosphoramidate before the measurement.

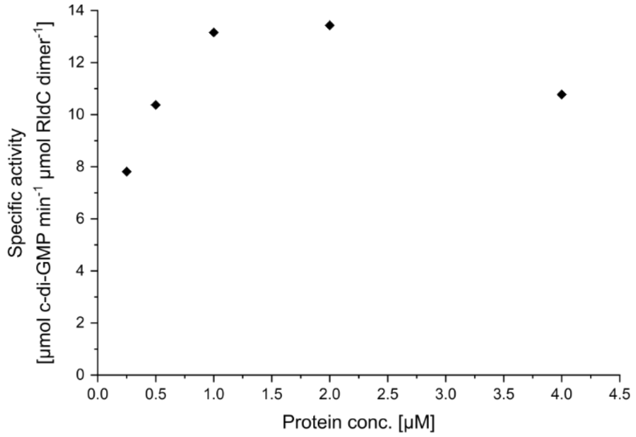


**Supplemental Figure 12: Concentration dependence of specific activity for *La*RldC.** The data indicates that lower concentrations than 1 µM show lower specific activity. This suggests the possibility of a monomer-dimer equilibrium influencing the protein's activity. Concentrations above 2 µM are affected by high substrate turnover, thereby limiting the achievable specific activity.

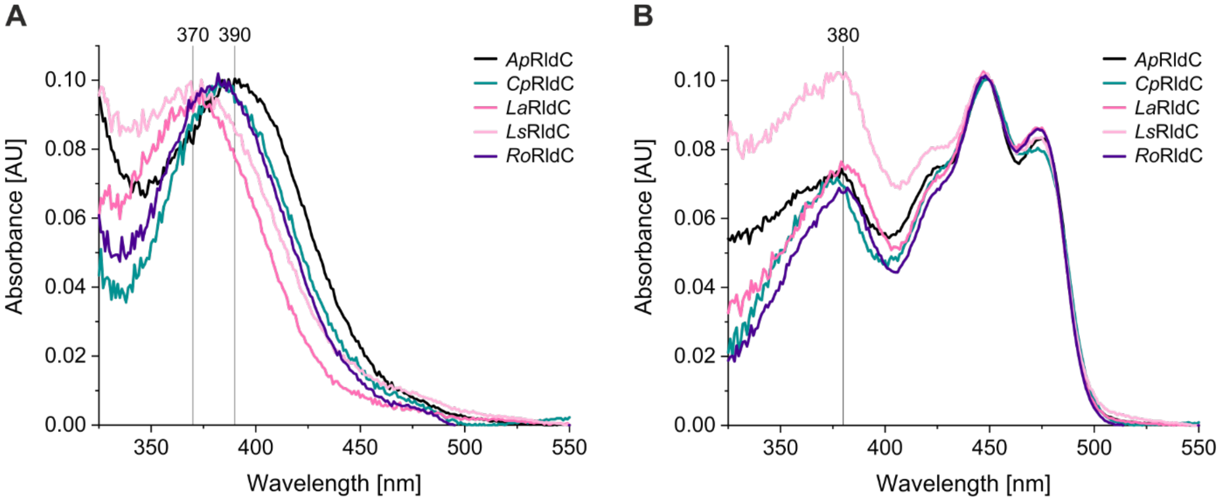


**Supplemental Figure 13: Differences in dark- and light-state absorption spectra of the studied RldC homologs.** A) The light-state of *La*RldC and *Ls*RldC is significantly blue-shifted compared to the usual 390 nm maximum observed in other LOV domains. Spectra normalised based on the observed maximum. B) Normalised dark-state spectra of all soluble homologs, comparing the maximum of the UV-A band.

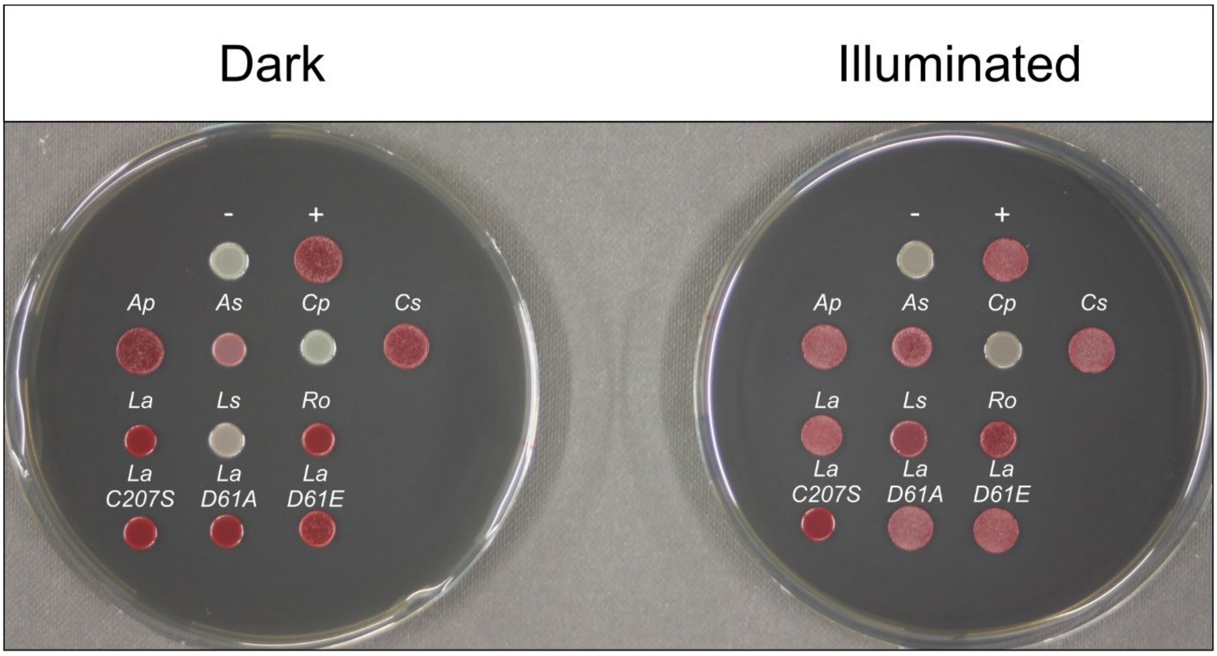


**Supplemental Figure 14: Uncut image of the in vivo screening plates shown in Figure 1.**

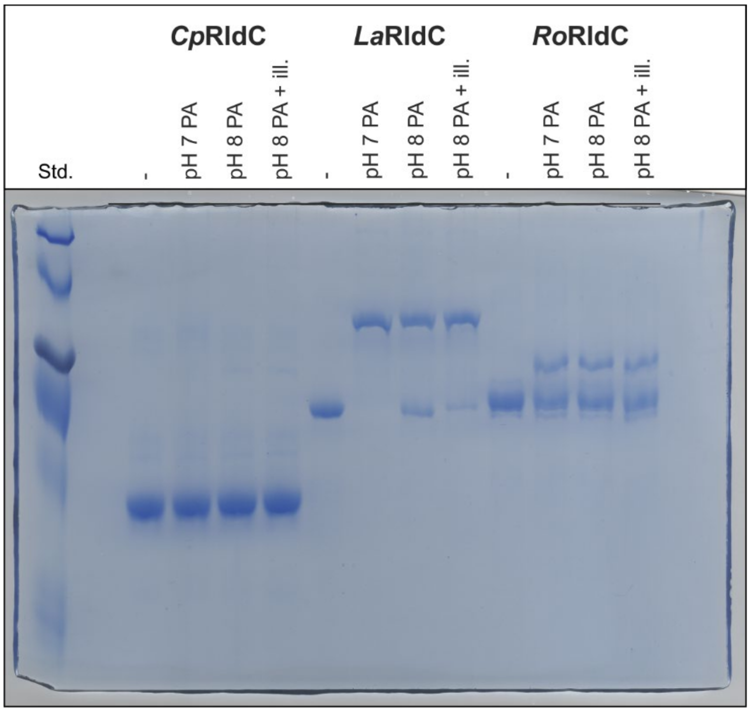


**Supplemental Figure 15: Complete Phos-Tag™ SDS-PAGE shown in Figure 5.** The coloured protein standard was used to follow the progress of the gel once the running front left the gel. Due to the gel composition, the protein standard cannot be used to estimate molecular masses.

**Supplemental Table 1: Data obtained from mass spectrometry measurements.** Expected masses were calculated using Prot pi (www.protpi.ch/Calculator/ProteinTool). Holo dimer masses contain the average masses of 2 molecules of FMN in addition to two protein protomers. The measurement accuracy was calculated in ppm.

| | Native MS | | | Intact Mass | | |
|---|---|---|---|---|---|---|
| **Protein** | **Expected Mass (holo dimer) [Da]** | **Measured Mass [Da]** | **Accuracy [ppm]** | **Expected Mass [Da]** | **Measured Mass [Da]** | **Accuracy [ppm]** |
| ***La*RldC** | 108,744.3 | 108,747.0 | 24.8 | 53,915.8 | 53,915.7 | -1.9 |
| ***Ls*RldC** | 115,373.5 | 115,376.5 | 26.0 | 57,230.4 | 57,230.3 | -1.7 |
| ***Ap*RldC** | 114,164.1 | 114,165.3 | 10.5 | 56,625.7 | 56,623.2 | -44.2 |
| ***Cp*RldC** | 108,200.0 | 108,201.6 | 14.8 | 53,643.7 | 53,643.1 | -11.2 |
| ***Ro*RldC** | 110,761.1 | 110,763.2 | 19.0 | 54,924.2 | 54,923.6 | -10.9 |

**Supplemental Table 2: Storage buffer composition for each of the studied homologs.**

| | Buffer composition | | | |
|---|---|---|---|---|
| ***La*RldC** | 10 mM Tris pH 8.0 | 500 mM NaCl | 2 mM $MgCl_2$ | 1 mM TCEP |
| ***Ls*RldC** | 10 mM Tris pH 8.0 | 500 mM NaCl | 2 mM $MgCl_2$ | 1 mM TCEP |
| ***Ap*RldC** | 10 mM Tris pH 8.0 | 50 mM NaCl | 2 mM $MgCl_2$ | |
| ***Cp*RldC** | 10 mM Hepes pH 7.0 | 500 mM NaCl | 2 mM $MgCl_2$ | |
| ***Ro*RldC** | 10 mM Tris pH 8.0 | 500 mM NaCl | 2 mM $MgCl_2$ | |

### Supplemental References

[77] Wolf D. Lehmann, Ralf Krüger, Mogjiborahman Salek, Chien-Wen Hung, Florian Wolschin, Wolfram Weckwerth, "Neutral loss-based phosphopeptide recognition: a collection of caveats", Journal of proteome research, Vol. 6, No. 7, pp. 2866–2873, 2007.
[78] Li Cui, Ipek Yapici, Babak Borhan, Gavin E. Reid, "Quantification of competing H3PO4 versus HPO3 + H2O neutral losses from regioselective 18O-labeled phosphopeptides", Journal of the American Society for Mass Spectrometry, Vol. 25, No. 1, pp. 141–148, 2014.
